# What are all these soil fungi doing? Complex carbon use ability as a predictive trait for fungal community functions

**DOI:** 10.1101/2025.07.11.664336

**Authors:** Tessa Camenzind, Denise Vonhoegen, Abdelhady Elshal, Rebeca Leme Oliva, Liam Whitehead, Corry Martin, Stefan Hempel, Christoph C Tebbe, Matthias C Rillig, Sören Thiele-Bruhn, Damien Finn

**Author notes:** Corresponding author: Tessa Camenzind – address: Altensteinstr. 6, 14195 Berlin, Germany; mail.

## Abstract

Fungal communities in soil play important roles in decomposition processes and soil organic carbon cycling. These communities are tremendously diverse, making it challenging to assign relevant functions to individual species. Fungal communities may be differentiated at the level of functional guilds; beyond such broad classification we have little delimitation, especially in fungal taxa common in grassland and agricultural soils. To resolve the level of functional similarity in fungal communities and define traits predictive of soil carbon cycling, we characterized fungal isolates abundant in agricultural soils to test the hypotheses that (i) the majority of saprobic soil fungi have the ability to use complex carbon sources, (ii) differences in complex carbon use abilities correlate with fungal enzymatic profiles, following principles of the fungal economics spectrum and (iii) carbon use ability is a predictive trait for fungal community functions. Using specialized growth media, we isolated and characterized 105 isolates and developed a novel *FungiResp* approach that directly tests fungal activity on complex carbon sources. The largest amount of variance between isolates was explained by differential abilities to use cellulose and starch, with only few phylogenetically distinct fungal clades showing high respiratory activity on these biopolymers. A preference for bacterial necromass was another major distinction among taxa. These key traits correlated with soil fungal community shifts in response to carbon substrate availability. By contrast, enzymatic activity was a poor predictor of fungal carbon use ability, except for correlations in lignin use and laccase activity. The newly established functional trait of carbon use ability offers important insights into diverse fungal communities: Many taxa lack the ability to use complex carbon (on their own), while the most common enzymes analyzed in soil showed little correlation with fungal mineralization potential. The discovery of key functional traits is an important step towards predicting the significance of fungal community shifts for soil carbon cycling.

## Introduction

Soils harbour tremendous phylogenetic and functional microbial diversity with crucial roles for soil processes. Though tiny in size as an individual, as a community microorganisms represent primary drivers of soil carbon and nutrient cycling. The breakdown of complex plant residues is an especially important ecosystem process determined by soil microbial activity. Predicting changes in mineralization dynamics in response to microbial community shifts remains challenging though (Yang 2021). Extensive knowledge of microbial community composition in response to global change drivers or agricultural management is available, but we still lack functional information on community members to make inferences for soil carbon cycling. Trait-based approaches in microbiology are in their infancy, lacking a broad coverage of taxa and in depth exploration of predictive traits (Lajoie and Kembel 2019). Total microbial abundance and potential activity may be estimated in soil samples – i.e., enzymatic activity, microbial biomass estimates or respiration (Joergensen and Wichern 2008). However, microbial biomass in soil is generally related to more easily available resources, whereas the breakdown of complex carbon is expected to be driven by a few taxa with high enzymatic capacities. Especially saprobic fungal species are known for their capacity to mineralize complex biopolymers like cellulose and lignin (Algora Gallardo et al. 2021). Most of such species specific information is available for forest-based wood degrading fungi, while common taxa in non-woody agricultural or grassland systems are less understood.

To interpret soil microbial community dynamics among different environments and conditions, the phylogenetic and functional resolution needed to distinguish microbial strategies must be resolved (Lennon et al. 2024). For soil fungi, classifications of taxa to functional guilds can be easily implemented with available trait databases (Nguyen et al. 2016, Põlme et al. 2020), though classification schemes are broad and often overlap (i.e., pathotrophs, saprotrophs and/or symbiotrophs). In fact, many soil fungi may even switch between different resource niches along a “saprotrophic-symbiotic continuum” (Aguilar-Trigueros et al. 2014), with their individual contribution to mineralization processes in soil being unclear. Generally, fungal carbon use ability is primarily researched in forest floor/wood fungal communities, dominated by white-rot and brown-rot Basidiomycete fungal groups differentiated based on wood-degrading capacities. By contrast, fungi abundant in agricultural and grassland soils are dominated by Ascomycota, Mucoromycota and Mortierellomycota (often referred to as “microfungi” or “moulds”), which are less well described in their enzymatic potential. Baldrian et al. (2011) report that only half of such non-basidiomycete isolates (15 out of 29 isolates tested) were able to degrade cellulose, also with low activity, and only 6 isolates showed laccase activity (phenoloxidase used for breakdown of lignin). The question remains to which degree the broad diversity of “other” soil fungi contributes to soil organic matter transformations in soil.

Individual fungal taxa show distinct functional capacities in carbon mineralization processes as shown in diverse experimental setups. In soil litterbag and successional litter decomposition studies, the availability of different biopolymers led to the formation of highly specific fungal communities (Hanson et al. 2008, Algora Gallardo et al. 2021; both studies focusing on forest ecosystem), with phylogenetically distinct groups corresponding to different carbon substrates like pectin, glucan, cellulose or chitin (Algora Gallardo et al. 2021). While such field studies give important insights on fungal carbon preferences in soil, some of these observations may be biased by underlying community dynamics. For example, the late occurrence of *Mortierella* species in litter succession is unlikely an indication of complex carbon preference (Vivelo and Bhatnagar 2019), but rather a consequence of specialization to microbial necromass utilization (Algora Gallardo et al. 2021, Morrissey et al. 2023). Thus, direct measurements under controlled laboratory conditions still represent an important tool to understand fungal carbon use ability. Laboratory experiments likewise have methodological limitations; not only is the number of isolates in culture limited, but also the degree to which such cultured isolates represent the full scale of potential traits in soil fungal communities. Fungal isolation techniques are biased towards taxa with simple sugar preferences (Hoefnagels 2005). Likewise, experimental laboratory conditions in agar media do not allow testing for complex carbon substrate use, since agar itself is a carbon source for fungi. Systems like the BioLog/FungiLog provide an interesting path forward to testing fungal substrate preferences in laboratory assays; however, the method also accounts for monomer use only (Singh 2009, Pawłowska et al. 2019). Polymer use may be additionally tested on agar plates, applying colorimetric measurement of a substrate hydrolysis zone (Baldrian et al. 2011, Pawłowska et al. 2019). Even though this approach gives insights into the ability of fungi to hydrolyze polymeric molecules, it impedes quantitative comparisons among different carbon substrates. As another important tool in modern ecology, *omics* may inform about fungal enzyme potential (Rosling et al. 2024). At a broader level of functional guilds, CAZyme (carbohydrate active enzyme) profiles revealed a gradual loss of plant cell wall degrading enzymes from saprobic to mycorrhizal lifestyles (Lebreton et al. 2021), while within the distinct classifications of white-, brown- and soft-rot fungi a wide functional CAZyme diversity was discovered (Floudas et al. 2020). For a more explicit application of these tools in the future, translation of genomic to phenotypic traits is needed based on an in depth understanding of functional traits predictive of carbon mineralization processes (Westoby et al. 2021, Li et al. 2023).

Enzymatic capacity can be easily measured in fungal isolates, and is commonly used as an important predictor of fungal carbon mineralization potential. However, due to the diversity of enzyme complexes involved in biopolymer degradation, predictions based on single enzyme assays (or also CAZyme families) may be insufficient (Li et al. 2023). Especially in regards to cellulose degradation - the most abundant carbon compound in (non-woody) plant residues - degradation involves complex interactions of endoglucanases, cellobiohydrolases and beta-glucosidases (Andlar et al. 2018, Moore et al. 2021), with some taxa showing even further mechanisms like Fenton-reactions (Floudas et al. 2020). To interpret enzyme assays, it is also important to note that enzymatic expression is regulated by the availability of simple versus complex carbon substrates, and is consequently reduced under classic culture conditions (i.e., glucose based media; Adnan et al. 2018). Most enzyme assays target individual enzymes involved in the degradation of specific biopolymers, whereas in soil a diverse mix of carbon compounds is present in soil organic matter. Specifically, non-forest soil systems may not be dominated by cellulose or lignin compounds, but include heterogeneous litter material from leaves and roots along with microbial residues (Gunina and Kuzyakov 2015, Liang et al. 2019). The mineralization potential of mixed substrates may not directly correlate with enzymatic profiles; not only does litter include diverse compound classes, but also priming is an important mechanism due to the presence of more labile carbon (Nottingham et al. 2009). Likewise, our knowledge of the degradation of microbial necromass as a relevant pathway of soil carbon cycling is incomplete. Studies either focus on individual chemical fractions of microbial cell walls (i.e., chitin or peptidoglycan) or fresh (autoclaved) microbial biomass that may not resemble more recalcitrant microbial residues that have undergone chemically transformative microbial death pathways (Camenzind et al. 2023). Since there are no individual enzymes predictive of mineralization of such heterogeneous carbon sources, direct substrate inoculation may more precisely reveal the level of substrate specialization in saprobic fungi (Algora Gallardo et al. 2021).

Insights into the ability of saprobic fungi to use complex carbon sources is of high ecological relevance. Improved functional trait information on carbon use dynamics is needed to predict resource niches of different fungal species, but also to link their life-history strategies with successional dynamics and responses to anthropogenic stressors (Camenzind et al. 2024). Life-history theory describes trade-offs in energy investment towards specific functions. Trade-offs in carbon mineralization and stress tolerance are especially important for predicting functional community shifts in response to environmental change or certain agricultural management techniques (Malik et al. 2020). Currently applied life-history theories in microorganisms have been derived from theoretical assumptions and macro-ecological findings, for example, the Y-A-S (yield – acquisition – stress tolerance) framework (Grime 1977, Malik et al. 2020). While such distinct life-history frameworks are useful for theoretical implementations of microbial traits into soil carbon models, postulated trade-offs do not have clear experimental support in saprobic fungi (Lustenhouwer et al. 2020, Alster et al. 2021a, Alster et al. 2022). The recently introduced fungal economics spectrum has also not supported trade-offs among Y-A-S categories; instead, complex carbon use ability and litter mineralization were orthogonal/linearly uncorrelated to growth rates and stress tolerance (Camenzind et al. 2024). Considering such lack of predictive trade-offs in carbon mineralization traits, novel theoretical frameworks must be explored for soil fungi. Morrissey et al. (2023) suggest analysing microbial traits relevant for soil carbon cycling in terms of consumers of (i) plant detritus, (ii) microbial necromass and (iii) dissolved organic carbon. In this view, fungal communities would be functionally characterized by their niche differentiation for specific resources. Since carbohydrate use ability (resource niche) is conserved at higher phylogenetic levels in fungi (Treseder and Lennon 2015, Rosling et al. 2024), phylogenetic trait imputation would further allow to functionally characterize diverse soil microbial communities based on a limited subset of studied isolates - even given the uncertainties attached to this method (Goberna and Verdú 2016, Karaoz and Brodie 2022). In order to develop such carbon cycling trait frameworks, we need further in-depth analyses of traits related to the resource niche in fungi, as well as their integration into classical life-history frameworks.

In this study we aim for an ecological understanding of complex carbon use ability as a functional trait in soil saprobic fungi. We tested 105 fungal isolates common in agricultural soils of Germany for their ability to use different carbon substrates, and evaluated the predictive power of this trait for soil community shifts (Fig. 1). Classical fungal traits were characterized, including characteristics of the fungal economics spectrum (Camenzind et al. 2024) as well as enzymatic activity profiles in the presence of simple vs. complex carbon substrates. To directly assess complex carbon use ability by fungi, we developed a novel *FungiResp* approach based on principles of BioLog and MicroResp analyses, which directly allows monitoring fungal activity on a wide array of carbon substrates. Resulting traits were used for phylogeny-based trait predictions, testing the predictive power of these traits for functional shifts in soil fungal communities. This large dataset allowed us to test the hypotheses that (i) the majority of soil fungi characterized as saprobes have the ability to use complex, but distinct carbon sources, (ii) differences in complex carbon use abilities may be predicted based on enzymatic profiles conserved at higher phylogenetic levels, rather than correlations with the fungal economics spectrum and (iii) carbon use ability is a predictive trait to interpret fungal community shifts.

**Fig. 1:**
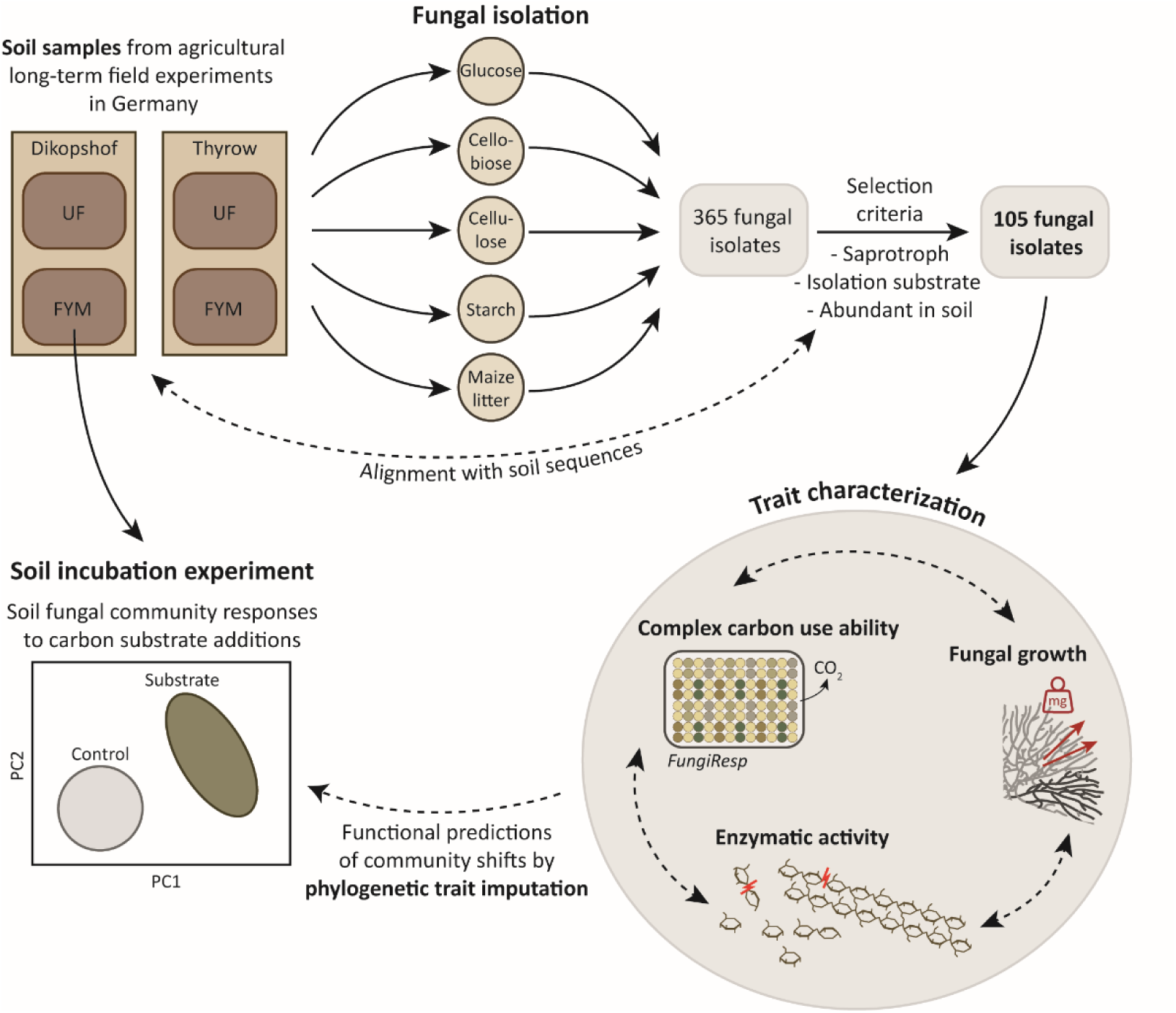
Overview graph of the complete experimental design, visualizing experimental steps (solid lines) and statistical analyses applied to combine different datasets (dashed lines). Dashed lines between different trait categories indicate correlations and trade-offs tested among functional traits. UF: unfertilized soils; FYM: Soils amended with farmyard manure.

## Materials and methods

### Fungal isolate collection

Fungi were isolated from German agricultural soils sampled in September 2021, including two sites with two fertilization regimes each (hereafter referred to as four soil types). Dikopshof and Thyrow are long-term experimental sites by the University of Bonn and Humboldt-University of Berlin, respectively. At each site, we chose plots fertilized with farmyard manure and unfertilized control plots. For detailed characterization and soil properties of these sites see Table S1 (more details are given by Ellmer and Baumecker 2002, Holthusen et al. 2012). Classical fungal isolation techniques were modified to target saprobic fungi with the ability to use diverse carbon substrates, which are active and abundant in the respective soils. Isolation included the following steps: (1) Soil washing to remove spores in order to capture active hyphae only, (2) dilution plating on simple and complex carbon sources, (3) individualization and identification of fungal isolates and (4) selection of fungal isolates based on their presence/abundance in soil and functional classification into saprotrophic guilds.

(1) Soil samples (top soils sampled in September 2021) from individual sites and treatments (stored at 4°C for 2-4 weeks) were mixed well and sieved to 2 mm with a sterile sieve. For each soil type, 15 × 5g of soil was shaken with 50 mL of 0.1% sodium pyrophosphate solution for one hour. Thereafter, the solution was transferred to a sterile 53 µm mesh and washed with dist. H_2_O for one minute (Parkinson and Williams 1961). The material kept on the 53 µm sieve was captured in a flask with 20 mL sterile H_2_O, and further used for dilution plating on different carbon substrate media.

(2) Defined growth media designed in previous studies were used for fungal isolation (Table S2; Camenzind et al. 2020, Leifheit et al. 2024). Glucose and cellobiose were applied as defined resource media with phytagel, using petri dishes of 6 mm in diameter (Table S2). Since phytagel on its own may be used as a carbon source by fungi, cellulose, starch and maize litter substrates were added as a thin layer of powder to small petri dishes (Ø 6mm) and all nutrients except carbon were supplied as liquid medium (Table S2, Fig. S1; only enough liquid was added to wet the carbon substrate powders). 200 µL of all prepared soil solutions (15 solutions per soil type from step 1) were added to respective media. For cellulose, starch and litter media, full solutions and 10^-1^ dilutions were applied, on glucose and cellobiose media 10^-1^ and 10^-2^ dilutions were evenly spread to separate individual colonies for further isolation (Fig. S1).

(3) After 3-6 days of growth, individual colonies were picked and transferred first to a medium based on the same carbon source, this time using phytagel based media for all carbon substrates, and in a second step transferred to malt-extract agar (10 g L^-1^ malt extract, 0.93 g L^-1^ peptone (extracted from soy) and 20 g L^-1^ agar; C:N ∼ 20) for further culturing. As a result, we obtained 200 fungal isolates per soil type, which we further reduced to 100 individual isolates with distinct mycelial morphologies. For the selected 100 isolates per soil type, fungal DNA was extracted using the Soil DNA Purification Kit (Roboklon GmbH, Berlin, Germany), and fungal identity determined by Sanger-sequencing, generating long sequences (about 1000bp) of partial SSU, ITS and partial LSU regions with ITS1f (Gardes and Bruns 1993) and NL4 primers (Kurtzman and Robnett 1998). Individual marker regions were compared to respective databases to determine fungal identity: ITS regions to the Unite database (Nilsson et al. 2018) and LSU to RDP LSU dataset (Cole et al. 2014).

(4) In order to select for isolates dominating respective soils, we characterized fungal communities in original soil samples via amplicon sequencing. DNA was extracted from 500 mg of four replicate samples of each soil type via the FastDNA Spin Kit (MP Biomedicals, Eschwege, Germany) following the manufacturer’s recommendations. For fungal community analyses, partial fungal ITS1 sequences were amplified with barcoded ITS1f and ITS2 primers (Walters et al. 2015, Hoggard et al. 2018) in 25 µL PCR reactions using Q5 high-fidelity DNA polymerase (New England Biolabs, Ipswich, MA, USA) as described previously (Finn et al. 2023). Amplicon products were purified and normalized to 2 ng µL^-1^ prior to library prep with Ovation Rapid DR Multiplex System 1-96 (NuGEN Technologies, San Carlos, CA, USA) and sequencing on the Illumina MiSeq platform by LGC Genomics GmbH, Berlin, Germany). Sequences were analyzed using the DADA2 ITS Pipeline Workflow (1.8) (https://benjjneb.github.io/dada2/ITS_workflow.html; Callahan et al. 2016), and assigned to taxa using the UNITE database (Nilsson et al. 2018; Fig. S2). In order to visualize the relationship between fungal isolates and abundant soil taxa, operational taxonomic units (OTUs, defined at 97% similarity, rarefied to a minimum of 40000 sequences per sample based on rarefaction curves (*vegan*; Oksanen et al. 2022)) found in soil were aligned with respective fungal isolate ITS1 sequences (AlignSeqs(), *DECIPHER* (Wright 2016)), and a neighbour joining tree constructed with the most abundant taxa (*phangorn* (Schliep 2010), Fig. S3, S4). From each soil type, 24-28 fungal isolates were chosen (Table S3) based on the criteria to be (i) characterized as “Saprotroph” by the fungal trait database FunGuild (Table S3; Nguyen et al. 2016) or other literature sources, (ii) close proximity or overlap with abundant taxa in soil (Fig. S3, S4) and (iii) aiming for an even representation of carbon sources originally used for isolation (Fig. 2, S5, S6).

**Fig. 2:**
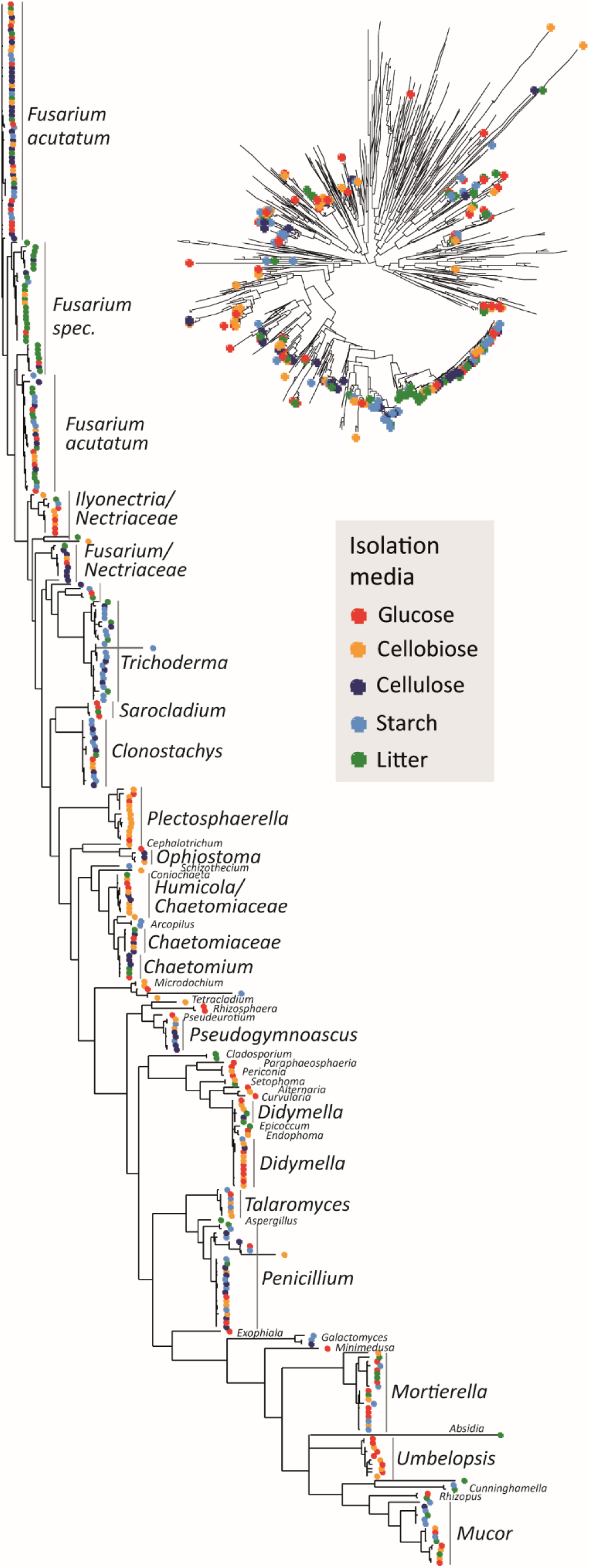
Phylogenetic tree depicting all fungal isolates obtained in this study by specialized isolation protocols on different carbon substrate media (indicated by colors). The radial tree displays phylogenetic placement of fungal isolates in relation to the 500 most abundant fungal taxa in all soils (Neighbour-joining tree based on ITS1 sequences). The phylogenetic tree depicting fungal isolates was built with long sequence reads (SSU, ITS, LSU region; Neighbor-joining tree based on MAFFT alignment). Identity of tips are visualized by genus labels, for a labelled tree with all isolate identities please refer to Fig. S6.

### Experimental design of trait analyses

All selected fungal isolates were tested for their ability to use different carbon substrates, applying a newly developed design that integrates principles of BioLog/FungiLog (Sobek and Zak 2003) and MicroResp analyses (Campbell et al. 2003), hereafter referred to as the *FungiResp* approach (Fig. S7). Experiments were run in 96 well plates for 7 days, adding carbon substrates in defined liquid media to quartz sand to create artificial soil conditions (Table S4). As an estimate of fungal activity, respiration was measured with MicroResp plates, and comparisons to control samples (no carbon source) were used to assess the ability of fungal isolates to use the respective carbon sources. Subsamples were further used to test enzymatic activity at the end of experiments. In parallel, further fungal traits representing the fungal economics spectrum (Camenzind et al. 2024) were characterized by classical agar plate designs, including mycelial extension rates, density and melanin contents. Correlations and phylogenetic signals in all phenotypic trait data for 105 fungal isolates were analysed in detail, and thereafter tested for their predictive ability for soil fungal community shifts in response to resource availability.

#### FungiResp design

Each well of 96-deep well plates was filled with 0.5 mL of furnace-dried quartz sand (particle size 0.1–0.5 mm; Sand-Schulz GmbH, Berlin, Germany) and autoclaved twice (Fig. S7a). Liquid media with different carbon sources were added (Table S4). Growth media were designed following principles according to Liebig’s Law of the Minimum (Camenzind et al. 2020): All elements were provided in sufficient amounts to ensure carbon to be the main limiting element and to observe clear effects of carbon substrate manipulation. Nutrient supply rates were determined based on the direct analysis of respective average element contents in fungal tissues, ensuring sufficient nutrient supply for fungal hyphal development while avoiding osmotic stress by high salt additions in conditions of reduced growth with complex carbon sources (Table S4; Camenzind et al. 2024, Leifheit et al. 2024). We prepared eleven different media with carbon substrates differing in complexity and heterogeneity: Glucose, cellobiose, maltotetraose, hemicellulose/xylan, cellulose, starch, lignin, litter, fungal necromass and bacterial necromass. Additionally, we supplemented apparently “recalcitrant” carbon sources with 5% (relative molar content) of glucose, to test for potential limitations of fungal carbon use by activation energy/priming (Highley 1977). Such glucose supplemented treatments are referred to as cellulose_act, starch_act and lignin_act (the ‘act’ standing for activation). Litter substrate was provided as maize leaves harvested at the end of the growing season. Maize plants were cultured at the Max-Planck-Institute for Biogeochemistry in Jena following conditions applied by IsoLife (www.isolife.nl, Wageningen, Netherlands; in collaboration with Marion Schrumpf). Fungal necromass was produced with a mixture of ten fungal isolates (DiF10, DiF19, DiF33, DiF35, DiF38, DiF56, DiF75, DiF78, DiF08, DiF94; Table S3), each cultured for six weeks on large glass petri dishes (Ø 20 cm) with malt-extract agar. Mycelial parts that were more than two weeks old were harvested to obtain realistic necromass (Camenzind et al. 2023). Similarly, nine bacterial strains (*Bacillus subtilis, Mycobacterium phlei, Micrococcus luteus, Pseudomonas fluorescens, Agrobacterium radiobacter, Arthrobacter globiformis, Nocardioides soli, Streptomyces griseus* and *Escherichia coli*) were grown and shaken in R2A liquid media for three weeks, and subsequently harvested by centrifugation and repeated washing steps of the bacterial pellets. Litter and evenly mixed fungal and bacterial necromass were freeze-dried and ball-milled. All carbon source additions were standardized by equal molar carbon additions (with a basis of 5 g L^-1^ glucose; Table S4). 180 µL of sterile medium was added to 96-well plate wells following a standardized design for all fungal isolates (Fig. S7a, b). Fifteen microliters of the liquid volume were used to apply fungal inoculum (see below), two replicates for each fungal isolate. Fungal respiration was monitored for 7 days applying the MicroResp plate approach (MicroResp™, James Hutton Ltd, Aberdeen, Scotland).

#### Fungal inoculum preparation

To avoid introducing carbon sources in the form of agar or extensive mycelial material, fungi were inoculated as defined colony-forming units (CFU) with sterile dist. H_2_O. One-sixteenth to one-eighth of fungal mycelia grown for 6 days on malt-extract agar plates (overlain with cellophane), cut and mixed from three replicate plates, were transferred to sterile Eppendorf tubes with water. The number of CFU were determined by transferring a defined amount of the solution (vortexed with metal balls) to malt-extract agar plates, counting CFU after 24 hours of incubation at 25°C. Fungal inoculum was applied as approximately 50 CFU in 15 µL solutions, with two replicates per treatment from the same fungal inoculum source for each isolate. This method allows for substrate colonization from spores and/or hyphal pieces, since fungal species show varying degrees of asexual sporulation in culture (Camenzind et al. 2022).

#### Respiration measurements

MicroResp CO_2_ detection plates were prepared aseptically following instructions given by the manufacturer (MicroResp^TM^). Autoclaved sealing mats were applied to the 96 well plates, and covered with cellophane between respiration measurements. From day two to seven of the experiment, detection plates were applied daily under sterile conditions for 4 hours, tightened by metal screw clamps. The absolute colour change in detection plates within 4 hours measured by a colorimetric microplate reader (570 nm) was used to estimate CO_2_ produced during this period of time. CO_2_ production [unitless] for each replicate, treatment and day was calculated by subtracting the mean respiration values in control treatments of respective isolates (n = 2; no carbon or control samples with 5% glucose, respectively). Thereafter, total CO_2_ production over the period of 7 days for each isolate and treatment was estimated as the area under the curve of respiration by time, calculating the integral function (generalized additive models, *mgcv*; Wood 2011). We tested in several runs for respiration activity during an extended period of 7 – 30 days, but no further increase in activity was recorded in any of the treatments or isolates tested. Sterile controls with glucose as a substrate were included to test for contamination during the experiment. There was no contamination, as there was no detectable respiration in control samples. A few replicates did not grow (displayed no activity) and were removed from further analyses.

#### Enzyme activity measurements

After the end of incubation, for a subset of 32 randomly selected isolates enzymatic activity was analyzed in a reduced amount of treatment combinations – glucose, hemicellulose, litter and cellulose_act, two replicates each. We used colorimetric assays to test for beta-glucosidase (substrate: 4-nitrophenyl-β-d-glucopyranosidase, duration of the reaction: 2 hrs), cellobiohydrolase (4-nitrophenyl-cellobioside, 4 hrs), beta-xylosidase (4-Nitrophenyl-β-D-xylopyranosid, 6 hrs) and laccase (2,2-azinobis-3-ethylbenzothiazoline-6-sulfonate (ABTS), 1 hr) activity. The upper 1 cm of sand/fungal mixture was transferred to Eppendorf tubes and mixed thoroughly. A defined volume (small spatula) was transferred to 96-well plates for each enzyme reaction, including a control without fungal substrate, and a control without enzyme substrate added. We added 150 µL sodium acetate buffer (pH 5) and 150 µL enzyme substrate at a concentration of 2 mM and incubated on a shaker (100 rpm) in the dark at 25°C for respective incubation periods. At the end of reactions, 100 µL of solutions were transferred to 96 well plates, and 50 µL 0.2 M NaOH added to end the reaction, except for laccase. Enzymatic activity was assessed by colorimetric changes measured at 410 nm in a microplate reader. To obtain relative enzymatic activity, colorimetric data measured for individual enzymes were subtracted from the maximum control value of the respective isolate and enzyme, using the maximum control value either of the enzyme control (no fungal material added) or the fungal control (no enzyme substrate added).

#### Fungal trait measurements on agar plates

Standard fungal growth traits were recorded following classic agar plate designs. Three replicates of each isolate were grown on malt-extract agar plates (Ø 9 cm). Mycelial extension rate [cm² day^-1^] was calculated based on mycelial size of individual replicates after three days of growth (using ImageJ software (Schneider et al. 2012)). Biomass [mg] and density [mg cm^-2^] of fungi were analyzed after 6 days of growth. Melanin contents were determined using a rapid colorimetric assay with Azure dye A (Fernandez and Koide 2014, Camenzind et al. 2024). We added 3 mg of freeze-dried and ball-milled fungal material to a tube, with 600 µL of Azure dye A in 0.1 M HCl (dye concentration adjusted to absorption of 0.67 at 610 nm). Following shaking head-over-head for 30 minutes, fungal material was deposited by centrifugation (8000 rpm for one minute). We transferred 100 µL of the solution to 96-well plates to measure absorption at 610 nm. Relative melanin concentrations are given as the absolute deviation among absorption values of fungal samples and the control (no fungal material added).

### Soil fungal community responses to substrate additions

In order to test for the predictive value of functional traits for soil fungal community shifts, we used sequencing data from a soil incubation experiment at the University of Trier. The incubation experiment was based on the same soil samples used for fungal isolation (described above) derived from the same soil sampling campaign in 2021. The incubation experiment was conducted with Dikopshof soil from the farmyard manure treatment (Table S1), which was sieved (< 2 mm) and air-dried. Prior to substrate addition, the soil was pre-incubated with 10 % [g/g] water to reactivate the soil. Starch and cellulose substrate isolated from *Zea mays* (IsoLife, Wageningen, Netherlands) were added, resulting in 1271 µg C g^-1^ soil dw (starch) and 1266 µg C g^-1^ soil dw (cellulose). The added substrate quantity is based on 800 % of microbial biomass carbon (1248 µg C g^-1^ soil dw), which was determined beforehand as sufficient to stimulate microbial activity. Microcosms (1 L Schott Bottles) were filled with a substrate-soil mixture in triplicates, accompanied by 3 controls (no amendment) per harvest time. A target bulk density of ∼1.15 g cm^-3^, maximum layer thickness of < 2 cm and water content of 16 % [g/g] (52% WHC) were applied. The microcosms were incubated at 20 ± 0.5 °C and destructively harvested after 4, 8, 16, 32 and 64 days, respectively. At these time points, subsamples were frozen and fungal communities analyzed via Illumina sequencing, following methods described above. Fungal abundance was measured as gene copies per gram dry weight soil via quantitative PCR using SYBR Green (ThermoFisher, Germany) as per the manufacturer’s instructions, targeting the ITS1 region, on a CFX96 Touch (Biorad Laboratories, Germany) system (Finn et al. 2023). Soil DNA solutions were standardized to 10 ng µL^-1^ prior to qPCR. *Fusarium culmorum* was used to generate standard curves (R^2^ > 0.99 and efficiency => 90%).

### Statistical analyses

All statistical and bioinformatic analyses were conducted in R version 4.4.1 (R Core Team 2021). The best phylogenetic tree of fungal isolates was obtained using alignments of long-read sequences with MAFFT (*ips*; Heibl 2008), creating a neighbor-joining tree (*phangorn*). Trees and plots were generated with *phyloseq* (McMurdie and Holmes 2013) and *ggtree* (Yu 2022). Phylogenetic distance was calculated as pairwise distances among sequences (*phangorn*) and visualized with non-metric multidimensional scaling (*vegan*). Effects of isolation media, soil type and soil fertilization treatments on phylogenetic patterns among fungal isolates (pairwise distances of sequences) were tested by permutational multivariate analysis of variance, PERMANOVA (*vegan*).

Phylogenetic signals in individual carbon substrate use traits were tested with *phyloSignal* (Keck et al. 2016) testing for Pageĺs λ. Phylogenetic correlations within the complete data matrix of carbon substrate use ability were tested by Mantel tests between Euclidean trait distance matrices and phylogenetic distances of isolates (*vegan*, *ape* (Paradis and Schliep 2018)). T-tests were applied to test for deviations among complex carbon use of cellulose, starch and lignin under primed versus non-primed conditions. To test for correlations with isolation media, carbon substrate use ability traits were visualized by principal component analyses (PCA; prcomp()), and shifts along the first two PC axes were tested by analyses of variances (ANOVA) of isolate PC scores in relation to original carbon isolation media.

Effects of carbon substrate, isolate identity and their interactions on enzymatic activity patterns were tested by PERMANOVA. Responses of the activity of individual enzymes to carbon substrate versus isolate identity were tested by generalized least square models (*nlme*; Pinheiro et al. 2021). This model controls for the heterogeneous variability among different fungal isolates, using log-transformed data to achieve normality.

Linear correlations among isolate traits were analyzed by PCA. To normalize the high variability in total respiration rates among isolates for trait correlations, carbon substrate use ability was standardized at a scale [0,1] relative to the maximum respiration rate observed within each isolate. Additionally, prior to PCA and further correlation analyses, all traits were tested for normality and log-transformed if necessary. The correlation structure of PCA was assessed by varimax rotation in case PC axes did not correlate well with individual traits (Carmona et al. 2021). To account for the impact of phylogenetic relatedness on trait correlations, we also performed phylogenetic PCAs using *lambda* as an optimization method (*phytools*; Revell 2012). Trait correlations among individual traits were calculated by Pearson correlation coefficients. Phylogenetically corrected linear correlations were fitted by phylogenetic generalized linear models (pgls(), *caper* (Orme et al. 2018)).

To test the predictive power of functional traits in soil, soil fungal OTUs assessed in the incubation experiment were characterized by phylogenetic trait imputation. Soil OTUs were aligned with fungal isolate ITS1 sequences using MAFFT (ips), and a Neighbor-joining tree was created. Combining the original trait matrix for isolates with the tree, functional traits were phylogenetically imputed to soil OTUs based on phylogenetic distances (*Rphylopars*; Goolsby et al. 2022). Using this method, we tested for functional shifts in carbon use ability of glucose, cellulose, starch, litter, bacterial and fungal necromass (relative values [0,1], see above), and the growth traits mycelial density, extension rate and melanin content.

Community-level weighted means of respective functional trait values were calculated based on Hellinger transformed community data (functcomp(), *FD* (Laliberté and Legendre 2010)). Correlations were plotted using unconstrained redundancy analyses of resulting community-weighted means of all traits assessed (*vegan*). Shifts in community-level weighted means in carbon addition treatments were further tested by analyses of variances, adding days as a within-group error term. Significant contrasts to control treatments were determined by estimated marginal means (*emmeans*; Lenth 2025). In order to interpret functional shifts in relation to characterized fungal isolates, we determined OTUs that significantly increased in response to carbon substrate additions using MaAslin 2, with treatment as a fixed factor and time as random factor, applying false discovery rate adjusted P-values (*Maaslin2*; Mallick et al. 2021).

## Results

### Fungal isolation

In total, we obtained 365 fungal isolates with long-read sequences from the four soil types (Fig. 2), from which we selected 105 fungal isolates for detailed trait analyses, evenly distributed by soil type and carbon substrate used for isolation (Fig. S3 and S4; Table S3). The isolation protocol with different carbon compounds did not lead to clearly distinct fungal isolates on respective media, in many cases isolates of the same species were derived from different carbon substrates (Fig. 2, S5, S6). Still, the phylogenetic distance among fungal isolates was significantly correlated with carbon substrate type (R² 0.10, *P* < 0.001; Fig. S5). In certain fungal genera/families we also observed a propensity for being isolated on either complex carbon media or simple glucose or cellobiose sources (Fig. 2). For example, species within the clade of *Trichoderma*, *Chaetomium*, *Pseudogymnoascus* or *Clonostachys* were preferentially isolated from cellulose or starch media. Whereas certain isolates affiliated with Nectriaceae, Didymellaceae, *Plectosphaerella*, Mortierellomycota or *Umbelopsis* (Mucoromycota) were primarily captured on cellobiose and glucose media, and never isolated on cellulose medium. Certain *Fusarium* species were again preferentially isolated on maize litter, though this clade was overall highly versatile.

We did not obtain significantly distinct fungal isolate collections from the two different soil sampling locations, only fertilization status of soil type was a significant (but weak) predictor of fungal isolate composition (R² 0.06, *P* = 0.02; Fig. S5). The low phylogenetic distinction between soil types was in contrast to the clear differentiation of fungal community composition observed among soils (Fig. S2). Interestingly, all soil types had a large proportion of Mortierellomycota (26 – 55% of total reads, Fig. S2b; compared to global abundance rates of this group below 10 % (Tedersoo et al. 2022)), which was also reflected in high numbers of *Mortierella* isolates (Fig. 2). Overall, we obtained a broad phylogenetic coverage of taxa present in soil, with abundant genera and taxa in soil also being represented in the isolate collection (Fig. 2, S3, S4); except for only one Basidiomycete (*Minimedusa spec*.), a phylum whose members are inherently challenging to isolate (Thorn et al. 1996). Though dominant in soil sequences (Fig. S3, S4), isolates of *Fusarium acutatum* were still overrepresented in the collection, and evenly isolated on all carbon media (Fig. 2). *Fusarium* species generally have high genetic and phenotypic plasticity, which potentially also biased the selection process: The strong variability in mycelial colors and shapes may have led to erroneous morphological selection of individual taxa.

**Fig. 3:**
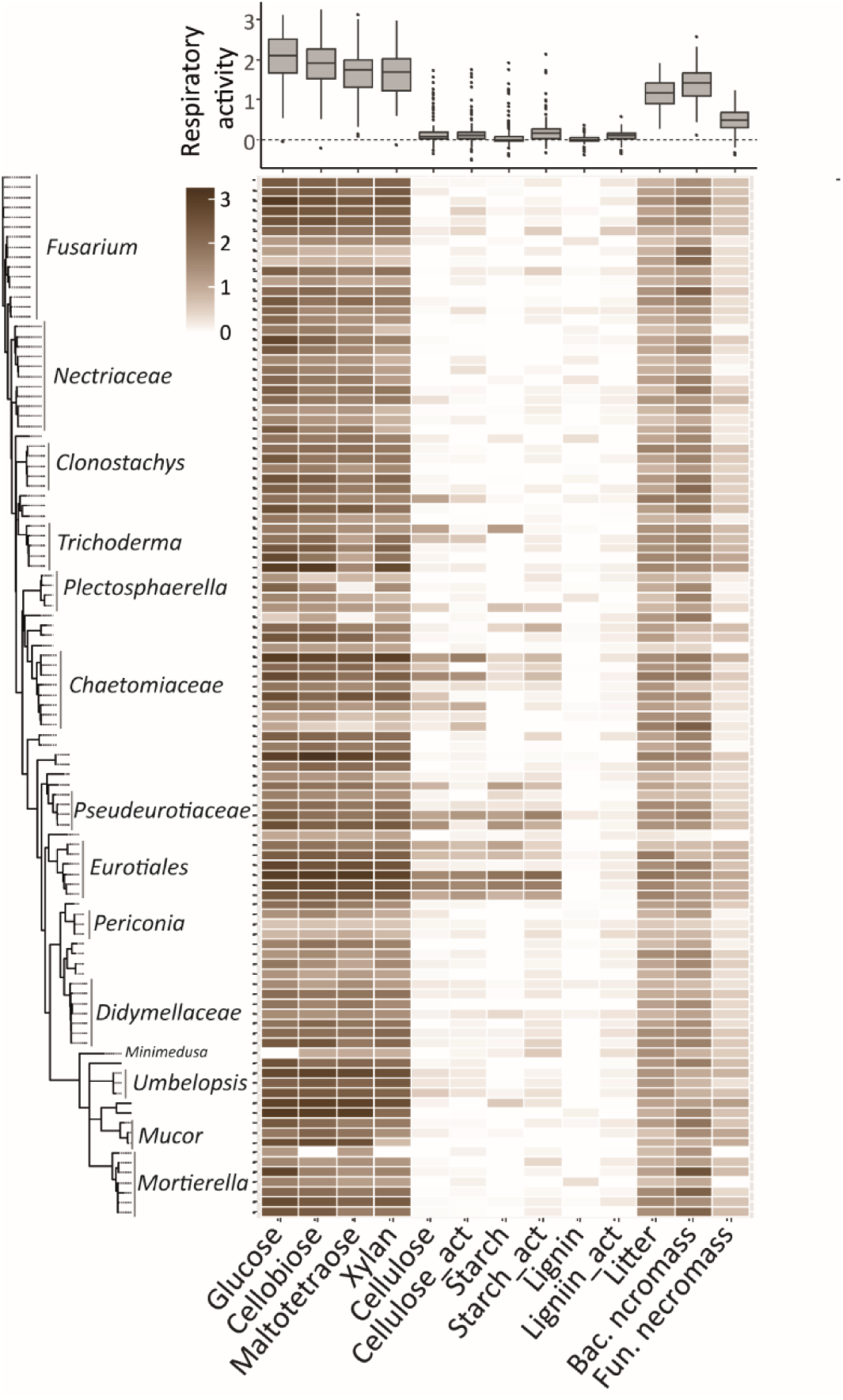
Carbon substrate use ability in diverse fungal isolates. The heatmap visualizes fungal activity on carbon substrates varying in complexity, given as CO_2_ production over 7 days (n = 2; values represent integral functions over time, based on MicroResp values subtracted by the control (no C added) of respective isolates). Overall data distribution among treatments is visualized by boxplots on top of the graph. Isolates are sorted based on their phylogenetic placement (neighbor-joining tree; Fig. 2; complete tree tip labels are given in Fig. S9).

### Ability to use complex carbon substrates

Fungal isolates showed high variability in their activity on different carbon substrate media, regarding absolute numbers as well as timing of respiratory activity (Fig. S7). Still, cumulative respiration values over 7 days were rather consistent, with highest activity on simple carbon sources, followed by intermediate activity on heterogeneous substrates like litter and bacterial necromass, and relatively low activity on fungal necromass as a carbon substrate (Fig. 3). By contrast, respiration on cellulose, starch and lignin media was only a fraction compared to that observed for simple carbon substrates. Only few isolates showed high activity on these complex carbon sources (Fig. 3, Fig S7g, h), primarily related to distinct phylogenetic clades within *Trichoderma*, Chaetomiaceae, Pseudeurotiaceae and Eurotiales (*Penicillium* and *Talaromyces*). In line with this observation, we detected strong phylogenetic signals in the fungal ability to use individual carbon sources (Pageĺs λ 0.92 – 1.20; *P* < 0.001), except for lignin use (Pageĺs λ 0.51, *P* = 0.01). When analyzing the complete trait matrix though, no significant correlation was observed between phylogenetic distances of fungal isolates and the functional trait space of carbon substrate use ability (*r* = 0.05, *P* = 0.24; Mantel test). Additionally, we did not detect an overall correlation of carbon use ability with isolation media (Fig. S8), even though the phylogenetic clades displaying high activity on complex carbon sources did partly match clades preferentially isolated on these sources (Fig. 2, 3).

When analysing the multivariate trait space, cellulose and starch use ability were the primary drivers of variation in the data, indicated by high loadings on the first principal component (PC) axis that explained 27% of variance (Fig. 4a). By contrast, high relative activity on bacterial necromass clearly differentiated certain isolates, and explained large variance within the dataset captured on the second PC axis (19% explained variance; Fig. 4a), showing trade-offs with activity on simple carbon substrates (glucose, cellobiose or xylan). The ability to use different complex carbon substrates under primed and non-primed conditions was significantly correlated (Fig. 4a, 6). Complex carbon substrates including 5% glucose to prime enzymatic activity (_act) led to an overall increase in the use of starch and lignin (*P* < 0.01), while cellulose use did not show a significant response (Fig. 3). These responses were highly isolate-specific, with some isolates even displaying reduced respiratory activity under primed conditions. Interestingly, PC analyses correcting for phylogenetic relatedness resulted in very similar correlation structure (Fig. S10a), though loadings of primed treatments (_act) were clustered more closely on the second PC axis.

**Fig. 4:**
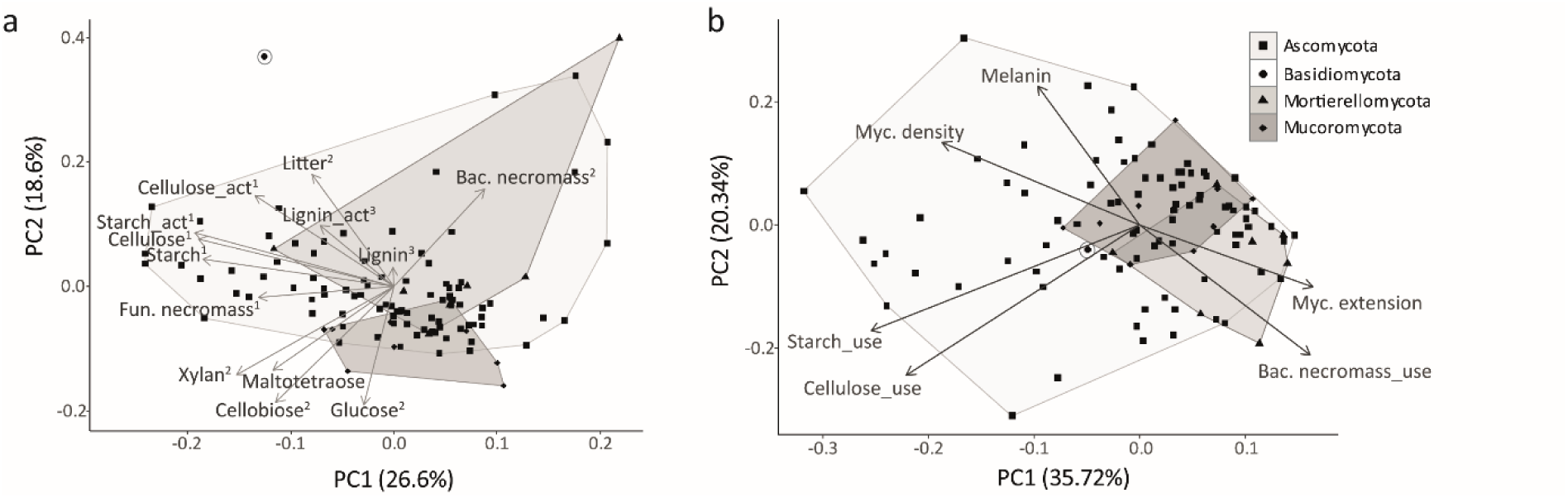
Principal component analyses (PCA) of the abilities of fungal isolates to use various carbon substrates (**a**) and correlations of selected carbon use traits to the fungal economics spectrum (**b**). Fungal carbon use ability on different substrates is given by relative values, standardized within individual isolates at a scale [0,1] relative to the maximum respiration rate. The first two principal component (PC) axes are displayed, including respective percentages of variability explained. Arrows represent eigenvectors of traits on PC axes, dots individual isolates (shapes of dots indicate phylogenetic placement). Shaded areas refer to the placement of phyla within the trait space. Superscript numbers (**a**) indicate traits with significant loadings on respective axes (*PCAtest*; Camargo 2022). Phylogenetically corrected PCAs are shown for comparison in Fig. S10. A varimax rotated PCA to clarify correlation structure (**b**) is displayed in Fig. S12.

### Enzymatic activity

Enzymatic activity measured at the end of the experimental period clearly varied among isolates and treatments (Fig. 5). Overall, isolate identity was strongly related to enzymatic activity patterns (R² 0.45, *P* < 0.001; PERMANOVA results), though carbon substrate effects (R² 0.15, *P* < 0.001) and especially the interactions of treatment and isolate identity (R² 0.34, *P* < 0.001) also were important predictors. Similarly, activities of individual enzymes were affected by isolate identity and carbon substrate type (Fig. 5). Carbon substrate type was a stronger predictor than isolate identity in the case of beta-glucosidase, cellobiohydrolase and beta-xylosidase activity, whereas laccase activity was rather isolate-specific. Notably, within individual isolates, enzymatic activity strongly varied in response to carbon substrate availability, with the direction of this variability being isolate-specific. Even laccase activity, which is a highly specialized enzyme only present in few fungal species, was restricted in some isolates to certain carbon substrate conditions (Fig. 5d).

**Fig. 5:**
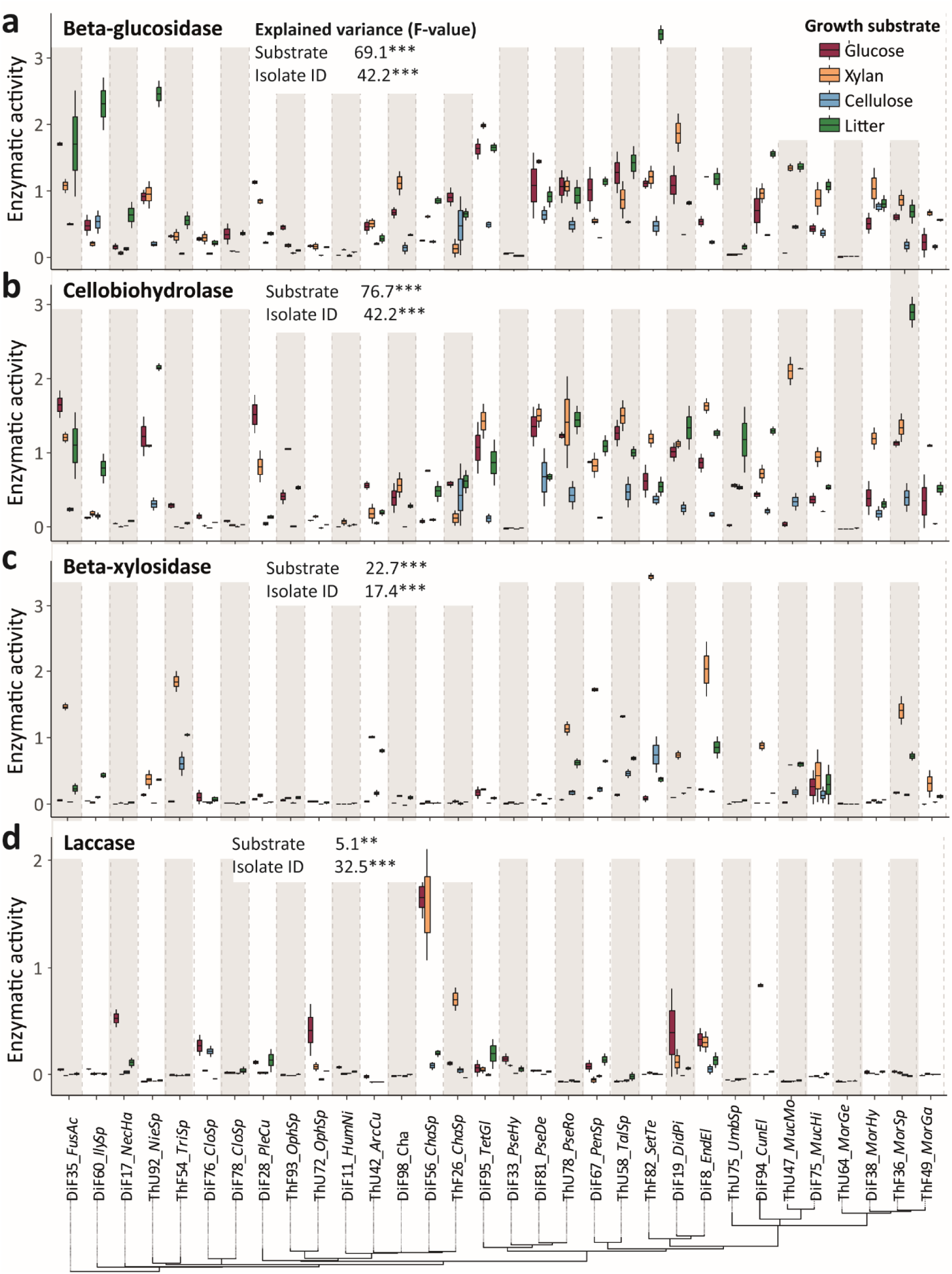
Enzymatic activity of fungal isolates grown on different carbon substrate media, tested for beta-glucosidase (**a**), cellobiohydrolase (**b**), beta-xylosidase (**c**) and laccase activity (**d**). Isolates are sorted based on phylogenetic placement, with labels referring to respective IDs and species abbreviations (Table S3). Carbon substrate treatments are visualized by colors of individual boxplots (n = 2). Effects of substrate vs. isolate identity on individual enzyme activities are given by respective F-values (generalized least square models; ** *P* < 0.01, *** *P* < 0.001).

Enzymatic activity levels were on average comparable among glucose, xylan and litter, and only slightly decreased in cellulose (Fig. S11). This finding is in contrast to fungal activity (respiration), which was reduced on complex carbon substrates in most isolates, with a sharp contrast especially between glucose and cellulose media (Fig. 3). Likewise, in contrast to enzymatic patterns, hyphal production was most pronounced on simple carbon substrates (visual observation). Consequently, in complex carbon media the proportional investment to enzyme production (in relation to fungal growth and activity) was increased compared to glucose media (Fig. S11).

### Correlation of functional traits related to carbon use

The ability to use different complex carbon substrates, especially of cellulose and starch, was correlated to each other (Fig. 4a, 6). Likewise, enzymatic activity of carbohydrate-active enzymes (except laccase) showed strong positive correlations with each other (Fig. 6). By contrast, enzymatic activity was not correlated with complex carbon use ability (Fig. 6), positive correlations were only observed for laccase activity and fungal activity on lignin media. Accounting for phylogenetic relatedness in linear correlations, xylan use revealed the expected positive relation to beta-xylosidase, as well as beta-glucosidase and cellobiohydrolase activity, while the preference for bacterial necromass was negatively correlated to these enzymes.

**Fig. 6:**
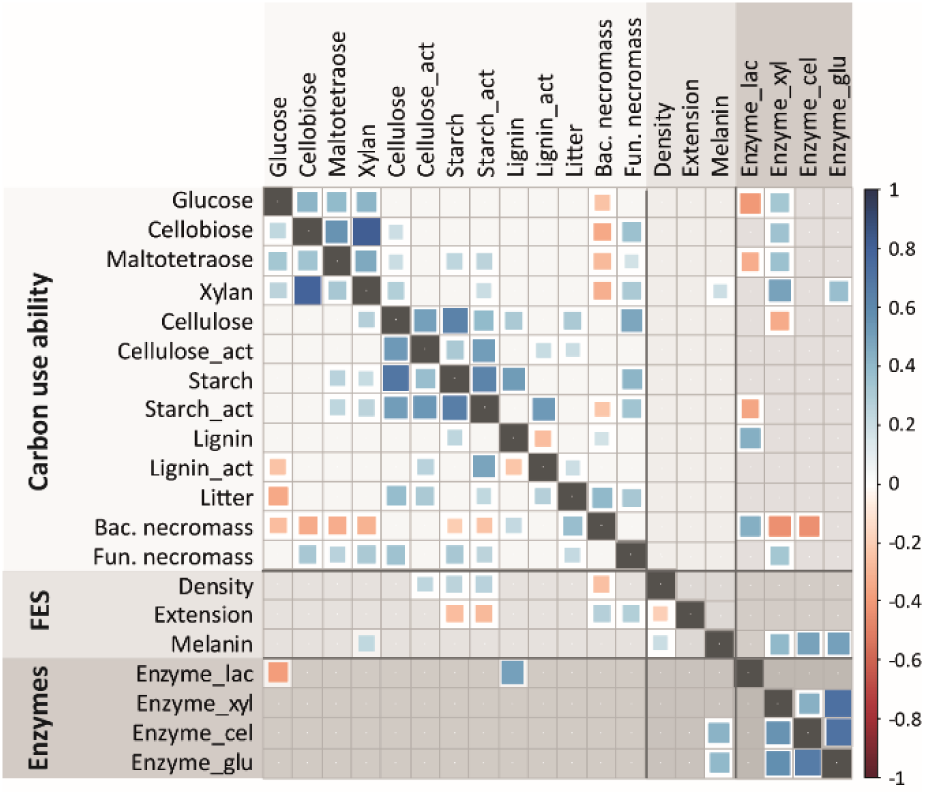
Correlation matrix of fungal isolate traits related to carbon use ability, the fungal economics spectrum (FES) and enzymatic activity. Pearson correlation coefficients are displayed on the lower left side of the graph, while the upper right side shows results of phylogenetically corrected linear correlations (square root of adjusted coefficients of determination; phylogenetic generalized linear models (Orme et al. 2018)); only significant values are displayed (*P* < 0.05). Enzymatic activity of individual enzymes (lac: laccase, xyl: beta-xylosidase, cel: cellobiohydrolase, glu: beta-glucosidase) were calculated for each isolate as mean values observed on different carbon substrate media.

Within the complete trait space, fungal ability to use complex carbon substrates was orthogonal (uncorrelated) to the fungal economics spectrum, here captured as the primary *dense-fast* trait continuum (Fig. 4b; Camenzind et al. 2024). As previously described in a smaller isolate collection (Camenzind et al. 2024), we observed a trade-off between slow growing fungi with dense mycelium and high melanin contents versus fast growing isolates with high mycelial extension rates (Fig. 4b, 6; correlations were partly driven by shared evolutionary history, Fig. 6, S10b). As previously described for the fungal economics space, loadings of complex carbohydrate use ability – cellulose and starch - were orthogonal to the dense-fast continuum (orthogonality was tested by varimax rotation of PC axes (Fig. S12)). When analysing individual trait correlations in more detail, functional traits related to slow growing fungi (high density, low mycelial extension rate) were positively associated with the ability of fungi to use complex C substrates, as shown by Pearson correlation coefficients (Fig. 6) and loadings on the first PC axis capturing most of the trait variation (Fig. 4b).

Similarly, melanin production (a costly cell wall component) was positively correlated to the overall fungal investment in enzyme production (Fig. 6). By contrast, fast mycelial growth rates were negatively correlated with substrate use ability of starch, but showed positive correlations with bacterial necromass use. It is important to note that some of these results were primarily driven by similar traits of closely related taxa, i.e., the correlation structure changed when correcting for phylogenetic relatedness (Fig. 6, S10b).

### Predictive traits for functional soil fungal community shifts

The composition of soil fungal communities shifted significantly in response to cellulose and starch additions in incubation experiments (R² 0.26, *P* < 0.001, PERMANOVA). Only cellulose additions led to a significant increase in fungal abundance measured by ITS copy numbers over time (on average 10 × 10^7^ ITS copies in cellulose amended soils after 64 days, compared to 3.8 × 10^7^ copies in starch treatments and 1.9 × 10^7^ in the control). Using phylogenetic imputation to assign functional traits, we also demonstrated a non-random shift in functional composition (Fig. 7a). Loadings of imputed functional traits (community-level weighted means) on PC axes showed an increase in the ability to use cellulose, starch and litter related to fungal communities in cellulose amended soils. These traits were significantly increased in cellulose treatments compared to the control (Fig. 7b). Likewise, fungal ability to use bacterial necromass showed an increase following cellulose additions (Fig. 7b). Mycelial extension rates and the ability to use fungal necromass were negatively correlated with cellulose additions. A range of OTUs showed significantly increased abundances in substrate amended soils (Fig. S13): Starch additions led to an increase in two OTUs related to Mortierellaceae, one Basidiomycete and some unclassified fungi. In cellulose treatments, primarily OTUs placed within Sordariomycetes, mostly Chaetomiaceae, were increased in abundance, as well as taxa within Eurotiomycetes (Gymnoascaceae) and Pseudeurotiaceae – all OTUs closely related to isolates characterized in this study with high complex carbon use ability (Fig. 3, Fig. S9).

**Fig. 7:**
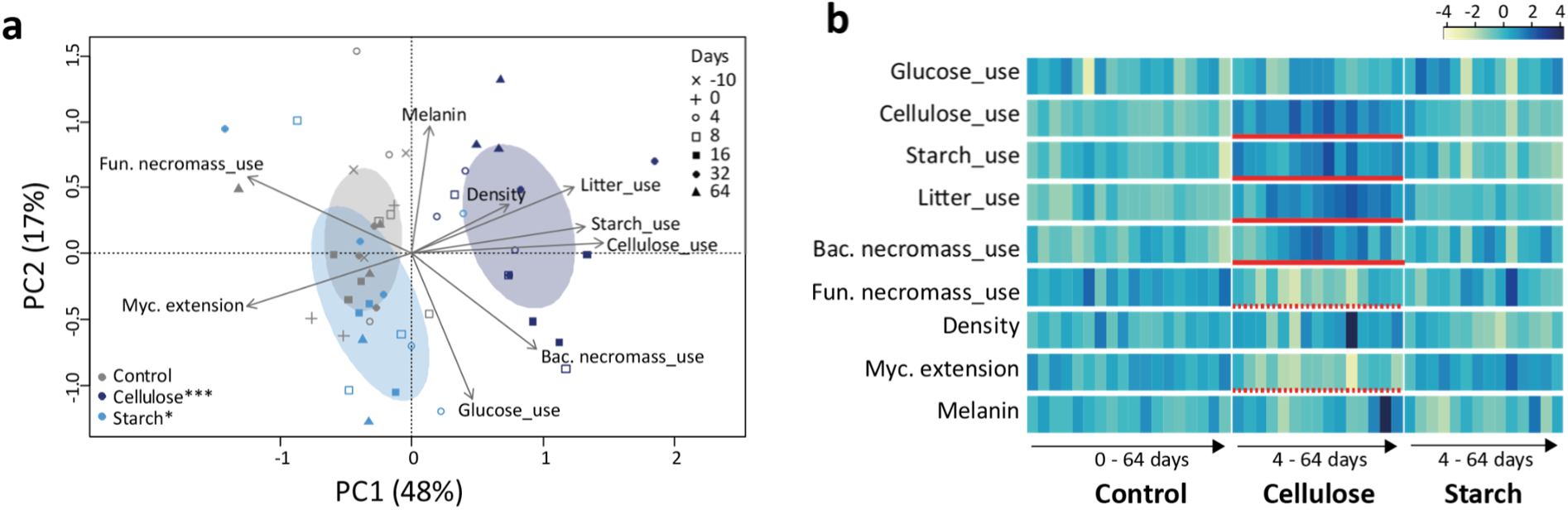
Functional soil fungal community shifts in response to carbon substrate additions, visualizing the multidimensional space of functional OTU composition (**a**) and shifts in community-level weighted means (**b**). **a** Principal component analysis (PCA) was performed to visualize the functional composition of soil fungal communities (trait matrix of community-level weighted means), characterized by phylogenetic trait imputation. Dots represent individual soil sample communities of different treatment (color) and sampling time (shape). Ellipses mark standard deviations of substrate treatments. Asterisks indicate significant differences in functional community composition in comparison to the control (pairwise permutational multivariate analysis of variance; * *P* < 0.05, *** *P* < 0.001). **b** Shifts in Community-level weighted means of individual functional traits are visualized as a heatmap, displaying functional shifts among treatments over time (n = 3). Darker colour indicates high relative values of scaled community-level weighted means (mean = 0, standard deviation = 1). Significant differences to the control are indicated by red lines, dashed red lines visualize negative effects.

## Discussion

We successfully isolated saprobic fungi broadly covering phylogenetic groups abundant in soils from two agricultural sites in Germany. When testing for the carbon use ability of the 105 selected isolates with our newly developed *FungiResp* approach, we found surprisingly wide variation especially in the capacity to mineralize cellulose and starch polymers. Despite the large diversity of saprobic fungi in soil, only a few fungal clades demonstrated an ability to degrade complex polymers as single isolates, contrary to our first hypothesis. We also partly reject our second hypothesis, since the activity of individual carbohydrate active enzymes was a poor predictor of these patterns: All tested fungal isolates showed activity of beta-glucosidase and cellobiohydrolase to varying degrees, and enzymatic activities were uncorrelated with cellulose and starch use ability. Only laccase showed specificity with isolates being able to use lignin. As hypothesized and previously described, complex carbon use ability was orthogonal to the main economics spectrum described for saprobic fungi, indicating only weak trade-offs in carbon mineralization traits with main life-history strategies. Instead, our data indicate that the ability to use complex carbon substrates represents a distinct niche dimension differentiating fungal clades. In support of our third hypothesis, we were able to use these traits to functionally characterize fungal community shifts in response to resource availability, a crucial first step to obtain functional insights into diverse soil fungal communities.

### Fungal isolation success on diverse carbon substrate media

Fungal isolation with different carbon substrate media led to a wide coverage of soil fungal communities, though only few fungal lineages showed high specificity to certain carbon substrates. Currently, the majority of fungal isolation efforts in soil are based on simple sugar media (Hoefnagels 2005), which limits functional trait characterization to fungi with likely reduced enzymatic capacities. The opposite extreme of fungal isolation produces isolates from mushrooms of wood decomposing fungi, characterized by high enzymatic potential (e.g., Lustenhouwer et al. 2020). Here, we aimed to broadly cover isolates along this gradient, which are abundant in agricultural non-woody systems. We did succeed in phylogenetically covering the dominant fungal taxa in these soils, as shown by isolate placements across the phylogenetic tree of soil OTUs. A preference of certain lineages to simple sugars, cellulose or litter substrate indicates that diverse carbon isolation media increased the coverage of soil fungal isolates. Despite this increase in coverage, isolation was still biased towards culturable taxa (Albright and Louca 2023). For example, despite clear differences in soil community composition among sites, comparable isolates were obtained from all soil types. Additionally, “blind spots” remain across the phylogenetic tree, concerning specific genera within Ascomycota, but also especially Basidiomycota and early-diverging fungal lineages (Fig. S3, S4; Thorn et al. 1996).

### Complex carbon use ability of soil saprobic fungi

Even though all of the selected isolates were characterized as potential “Saprotrophs” by FunGuild or other sources, surprisingly few clades showed significant activity on cellulose and starch. As expected, activity was highest on simple monomers, dimers or oligosaccharides of glucose, followed by the hemicellulose xylan, a soluble polymer of ∼80 – 200 xylose units. By contrast, fungal activity was strongly reduced in cellulose and starch polymers, and hardly detectable on lignin (in the absence of activation energy). Cellulose is the most abundant biopolymer in soil organic plant residues (Kögel-Knabner 2002), which is often discussed as a relatively labile carbon source for fungi, primarily in contrast to lignin. This view derives from a focus on enzymatic capacities in wood decomposing fungi, i.e., white-rot versus brown-rot and soft rot fungal groups (Eastwood et al. 2011). In agricultural soils, though, the majority of abundant fungal taxa appear to have limited ability to mineralize these polymers in isolation (our data; Baldrian et al. 2011). Cellulose consists of glucose assembled in chains with β-1,4-glycosidic bonds of hundreds to thousands units, which are further aligned in microfibrils via hydrogen bonds (Floudas et al. 2022). This complexity may explain the limited predictive value of beta-glucosidase and cellobiohydrolase activity, which are commonly tested enzymes for carbon mineralization dynamics in soil. Both enzymes only initiate the last steps of cellulose degradation to cellobiose and glucose units (exo-enzymes), whereas additional endo-enzymes are involved in preceding mineralization steps (Hsin et al. 2025). Baldrian et al. (2011) even report a negative correlation between endoglucanases and cellobiohydrolase activity in soil fungi, supporting the low predictive power of cellobiohydrolase for initial cellulose depolymerisation steps. It is further interesting to note that the degradation of cellulose and starch was highly correlated in soil fungal isolates, indicating either similar degradation steps of its crystalline form, despite different glycosidic bond types (Apriyanto et al. 2022), or a co-occurrence of respective enzyme complexes in fungal clades. Priming of polymer mineralization by glucose additions only lead to minor changes in fungal activity (Highley 1977), suggesting that low activity was a result of a lack of enzymatic capacities, rather than the absence of activation energy.

In mixed heterogeneous substrates fungal activity was relatively high compared to individual biopolymers, with few isolates showing distinct preferences for bacterial necromass. Bacterial necromass also resulted in the highest respiratory activity, followed by maize litter and comparatively low activities on fungal necromass. Fungal necromass used in our design was produced by natural senescence as a microbial death pathway (Camenzind et al. 2023), and likely primarily consisted of cell wall components. Since hyphal cell walls are primarily made of long chain glucans and mannans, chitin and melanin (Gow et al. 2017), fungal necromass is a rather recalcitrant carbon residue (in contrast to often used autoclaved biomass including more labile cytosolic compounds; See et al. 2021). Melanin contents may have reduced fungal necromass mineralization rates (Fernandez and Koide 2014), though only one of the ten isolates used for necromass production showed dark coloration. Nitrogen was no limiting factor in our experimental design, still, the molar C/N ratio of fungal necromass (molar C:N ratio = 37) in comparison to bacterial necromass (4.5) and maize litter (18) further indicates increased recalcitrance of the substrate. The ability of fungal isolates to use litter and fungal necromass were positively correlated with carbon use ability of cellulose and/or starch, suggesting a general ability to mineralize complex cell wall components. Whereas again, enzymatic activity of beta-glucosidase, xylosidase and cellobiohydrolase was also a poor predictor of heterogeneous carbon substrate use. Bacterial necromass gave rise to high mineralization rates in all fungal isolates, highlighting that it is a more labile carbon source as indicated by narrow C/N ratios (despite a similar approach of cell ageing/senescence as used to produce fungal necromass). The mix of Gram-positive and –negative cell residues used here likely results in more labile residues of membrane components, peptidoglycan and proteins (Silhavy et al. 2010). The distinct preference of few fungal isolates for bacterial necromass, indicated by high variance explained by this trait on the second principal component axis, supports the notion of resource specialization on microbial necromass residues as a strategy in microbial organisms (Morrissey et al. 2023).

### Integrating fungal carbon substrate niches in life-history strategies

In this dataset, the ability to use complex carbon sources (cellulose and starch) drove the main explanatory power of functional trait variation, and this ability did not strongly trade-off with other aspects of life-history strategies. In accordance with a previous trait collection in 28 grassland fungal isolates (Camenzind et al. 2024), complex carbon use ability was more positively related with dense/slow growing melanised fungi. Still, the complete trait space showed an orthogonal/uncorrelated structure between the main dense-fast economics spectrum and complex carbon use ability, which contradicts predicted trade-offs in acquisition and growth traits as applied for example in the Y-A-S framework (Malik et al. 2020). Instead, our data give support to the idea of niche-based microbial strategies (Morrissey et al. 2023), where niche preferences for labile vs. complex polymer resources are partly independent of primary life-history trade-offs related to growth. Only some of the tested carbon use traits may be found on the axis of the economics spectrum: Bacterial necromass use was positively correlated with fast growing fungal isolates. As observed for other correlation pairs, the positive correlation of bacterial necromass use with mycelial growth rates was primarily driven by phylogenetically related taxa (Fig. S10b), consequently it does not necessarily represent a universal functional trade-off but is rather driven by shared evolutionary history and trait selection (Agrawal and Rasmann 2010). In order to predict functional shifts in fungal communities related to carbon cycling, it is especially important to explore further trade-offs with stress tolerance and environmental predictors (Joswig et al. 2022). Fungal tolerance to moderate anthropogenic stressors has been associated with the fast side of the economics spectrum (Camenzind et al. 2024). If this correlation holds true, it would support low trade-offs in complex carbon use ability with stress tolerance (Jones et al. 2025).

Microbial life-history theory is currently under intense debate, with general challenges of trait ecology specifically applying to soil microbial traits that can rarely be measured within soil (de Bello et al. 2025). Several authors have recently advocated for a “bottom up” approach in microbial life-history theory, i.e., determine functional trade-offs in the organisms itself *a priori* rather than applying theoretical “top down” concepts (Romillac and Santorufo 2021, Westoby et al. 2021, Treseder 2023). While we agree with this endeavour, we here further emphasize the importance of selecting functional traits that are ecologically meaningful and that can be quantified in microbes. For example, mycelial growth rate was shown to be a meaningful isolate-specific trait: Mycelial growth rates measured on artificial agar media were correlated with hyphal growth dynamics in soil (Whalen et al. 2024) and demonstrated to be a robust isolate-specific trait under diverse environmental conditions (Camenzind et al. 2024). By contrast, enzymatic activity showed little isolate-specificity, with high plasticity depending on carbon substrate availability (Fig. 5). Also, the observed lack of correlation with actual carbon mineralization rates and substrate preferences calls into question its functional significance in an ecosystem context (Violle et al. 2007, Leifheit et al. 2024). Metabolic enzyme costs are an essential part of growth investment in saprobic organisms, but the way it is used as a functional trait needs methodological re-evaluation (Wortel et al. 2018, Gunina and Kuzyakov 2022). The lack of correlation in enzymatic activity and complex carbon use ability further challenges the interpretation of enzymatic assays in soil (commonly focusing on cellobiohydrolase and beta-glucosidase) as an indication of microbial carbon limitation and mineralization of plant polymers and soil organic matter (Moorhead et al. 2023). Testing a broader suite of enzymes (and their temporal dynamics) or adding direct tests of carbon substrate use may be more predictive of soil microbial mineralization potential.

### Predictions of functional shifts in soil fungal communities related to carbon substrate use

Phylogenetic imputation of trait values enabled us to characterize soil fungal communities, with functional shifts revealing ecologically significant responses to carbon substrate availability. Trait imputation methods were applicable due to phylogenetic signals detected in all tested functional traits; especially starch and cellulose use was conserved within specific fungal lineages (Fig. 3). Imputing unknown functions based on phylogenetic modelling is a common method in microbial trait research (Ruiz et al. 2023), often also applied for genetic traits with limited numbers of reference genomes (PICRUSt; Douglas et al. 2020). Still, this method comes with uncertainties (Goberna and Verdú 2016): (i) “blind spots” exist across the phylogenetic tree that are not covered by closely related taxa with known traits (Fig. 2) and (ii) high variability and plasticity in functional traits within species but also in closely related taxa add inaccuracies (Fig. 3; Alster et al. 2021b, Camenzind et al. 2024). In light of these uncertainties, the distinct non-random functional shifts observed here in soils amended with carbon substrates are astonishing. Especially cellulose addition, which led to increased fungal growth and distinct fungal communities, correlated with the predicted functional ability of fungal community members to use cellulose, starch and litter. With a slight time shift, also the relative abundance of OTUs characterized by preferential bacterial necromass use was increased; since soils were non-sterile, this may be an indirect effect of bacterial growth and death dynamics over time. Soil OTUs favoured by cellulose additions were indeed closely related to isolates characterized by high complex carbon use ability in our collection, i.e., members of Chaetomiaceae, Pseudeurotiaceae or Stachybotryaceae (Fig. S9, S13), also partly characterized as soft rot fungi (Darwish and Abdel-Azeem 2020). To differentiate these functional shifts from responses to simple carbon substrates, we analysed an experiment testing for maize litter and peptidoglycan additions in comparison (Fig. S14, unpublished data). In contrast to complex polymer effects, substrate additions induced most pronounced functional shifts in fungal preferences for glucose and bacterial necromass use (Fig. S14). In summary, despite large variability in the data and open research gaps, trait imputation revealed distinct and predictable functional shifts in the presence of complex vs. simple carbon substrates and proved a powerful method for functional soil community characterization.

Especially the trait of complex carbon use ability we established here represents an important step towards more precise functional characterizations of soil fungal communities. So far, databases like FunGuild or FungalTraits are used to characterize soil fungal strategies at a broader level (Nguyen et al. 2016, Põlme et al. 2020). These databases are useful to quantify plant-associated fungi, but provide little indication on carbon mineralization patterns of saprobic fungal communities in soil: Nearly all of the isolates tested here were classified as “Saprotrophs” (though many were additionally labelled as Symbiotrophs and/or Pathotrophs), of which only few were able to degrade complex biopolymers. None of the broader strategies given by FunGuild or FungalTraits were specific to tested carbon use traits (Fig. S15).

Similarly, FunGuild classification did not reveal significant functional shifts in soil fungal communities in response to substrate additions (data not shown). The advantage of these databases is their wide coverage of taxa, which is challenging to realize with more precise lab-based functional trait analyses. The increasing amount of genomic data may be key to achieve a similarly wide coverage with functional traits. As a first step, ecologically relevant genetic traits may be defined based on model predictions with phenotypic trait data (Morrison et al. 2022, Kijpornyongpan et al. 2025). If available, such data would allow functionally characterizing diverse fungal communities at global scale, similar to genetic predictions already widely applied in bacteria (Douglas et al. 2020, Karaoz and Brodie 2022). Such broad scale analyses also allow testing for trade-offs in carbon (resource) niche traits with environmental parameters and stress tolerance (Treseder et al. 2021).

Predicting community functions further requires accounting for the fact that these fungal lineages co-exist and interact in soil (de Bello et al. 2025). Contrary to common assumptions of primarily competitive interactions established for wood decomposers (Hiscox and Boddy 2017), positive interactions tend to prevail in soil microbial communities and litter inhabiting fungi (LeBauer 2010, Kost et al. 2023). Interaction mechanisms of cross-feeding and enzymatic complementarity may lead to more positive community interaction traits and increased carbon mineralization dynamics than predicted based on fungal traits in isolation (Geesink et al. 2024, Kaschper et al. in prep.).

## Conclusions

Soils harbour highly diverse fungal communities, which can be captured and described by high-throughput sequencing, while functional classification lags behind. Despite the high fungal diversity present in soil, we observed that only a limited number of distinct fungal lineages had the ability to mineralize complex biopolymers and cell wall components of plant and microbial residues. Consequently, only a few of the saprobic fungal taxa observed in soil may contribute to mineralization of polymeric carbon sources, an ability crucial for predicting decomposition rates and carbon release from soil (Schädel et al. 2013). Our data indicate that the majority of saprobic fungal taxa detected in agricultural soil sequences is primarily associated with labile plant or microbial residues, possibly also to root exudates in the rhizosphere or even living root tissues. Functional shifts in agricultural soil communities in response to substrate additions support this view: While cellulose availability favoured taxa closely related to the few lineages with high complex carbon use ability, more simple substrate additions of maize litter and peptidoglycan increased OTUs also predicted to have simple carbon preferences. These results demonstrate how the newly established fungal trait *complex carbon use ability* facilitates functional characterization of soil fungal communities, which is likely more powerful than predictions based on classical enzyme assays. Low correlations of complex carbon use ability with life-history strategies calls into question current assumptions of trade-offs between stress tolerance, fungal growth and acquisition traits (Finn et al. 2021). Future studies may connect these selected phenotypic traits with genomic data, in order to extend the coverage of fungal taxa and establish predictable trade-offs in carbon mineralization traits with environmental drivers. This opens up possibilities to connect the enormous amount of soil fungal community data with specific functions driving soil carbon cycling, making them predictable in biogeochemical models across environmental gradients.

## Acknowledgments

We thank Marion Schrumpf (Research group Soil Biogeochemistry, Max-Planck Institute for Biogeochemistry) for kindly providing the maize litter substrate. We are grateful to Annika Rüdinger and Irina Protasova (Freie Universität Berlin) for support during fungal culturing. TC acknowledges funding by the Deutsche Forschungsgemeinschaft (DFG, grant number 465123751, CA 1948/2-1, SPP2322 SoilSystems). SH acknowledges funding by the Deutsche Forschungsgemeinschaft (grant HE 6183/5-1). A part of this research was conducted within the DFG-project DriverPool (Th678/26-1, SPP2322 SoilSystems).

## Notes

### Competing Interest Statement

The authors have declared no competing interest.

