## Supporting Information S1 for "What are all these soil fungi doing? Complex carbon use ability as a predictive trait for fungal community functions"

**Journal:**

**Table S1** Agricultural sites sampled for fungal isolation.

| **Site** | | **Dikopshof** | | **Thyrow** | |
|---|---|---|---|---|---|
| **Treatment** | | **Farmyard manure** | **Unfertilized** | **Farmyard manure** | **Unfertilized** |
| **Location** | | 50°81′N, 6°95′E | | 52°16′N, 13°12′E | |
| **Annual mean temperature** | [°C] | 10.5 | | 9.2 | |
| **Annual mean rainfall** | [mm] | 688 | | 510 | |
| **Trial established** | | 1940 | | 1937 | |
| **Soil group** | | Haplic Luvisol | | Haplic Retisol | |
| **Texture** | | Silt loam | | Loamy sand | |
| TOC | [%] | 0.74 | 0.69 | 0.68 | 0.28 |
| TN | [%] | 0.08 | 0.08 | 0.06 | 0.03 |
| C/N | | 9.5 | 8.8 | 11.4 | 11.1 |
| pH | ($CaCl_2$) | 6.3 | 6.1 | 5 | 4.2 |
| P | [mg $kg^{-1}$] | 57 | 30 | 70 | 52 |
| K | [mg $kg^{-1}$] | 161 | 26 | 87 | 12 |
| Clay | [%] | 15.1 | 16.2 | 5.6 | 5.5 |
| Silt | [%] | 68.9 | 67.3 | 11.1 | 11.1 |
| Sand | [%] | 15.9 | 16.4 | 83.3 | 83.3 |

**Table S2** Defined growth media with different carbon sources used for fungal isolation (media were additionally supplemented with antibiotics to avoid bacterial growth (chlortetracycline, streptomycin and penicillin).

| **Element** | **Compound** | **Medium concentration** | **Comment** |
|---|---|---|---|
| C | glucose | 5 g $L^{-1}$ | Carbon was added at equal molar amounts; Carbon sources must be autoclaved separately to avoid medium |
| | cellobiose | 4.75 g $L^{-1}$ | |
| | cellulose | 4.4 g $L^{-1}$ | |

| | | | |
|---|---|---|---|
| | starch | 4.5 g $L^{-1}$ | caramelization;<br>pH values of carbon substrate solutions were adjusted to ~5.6 |
| | maize litter | 4.42 g $L^{-1}$ | |
| N | $NH_4NO_3$ | 0.503 g $L^{-1}$ | |
| P | $NaH_2PO_4$ | 0.183 g $L^{-1}$ | Phosphate must be autoclaved separately to avoid insoluble precipitates or toxic conditions |
| Mg, S | $MgSO_4$ | 0.134 g $L^{-1}$ | |
| K | KCl | 0.023 g $L^{-1}$ | |
| Ca | $CaCl_2$ | 0.065 g $L^{-1}$ | |
| Fe | NaFeEDTA | 0.02 g $L^{-1}$ | |
| Vitamine | Thiamine HCl | 1 mg $L^{-1}$ | |
| Vitamine | Biotin | 0.05 mg $L^{-1}$ | |
| Mo | $Na_2MoO_4$ | 0.05 mg $L^{-1}$ | |
| Cu | $CuSO_4$ | 0.01 mg $L^{-1}$ | |
| Mn | $MnSO_4$ | 0.05 mg $L^{-1}$ | |
| B | $H_3BO_3$ | 0.05 mg $L^{-1}$ | |
| Zn | $ZnSO_4$ | 5 mg $L^{-1}$ | |
| Gelling agent | Phytagel | 2 g $L^{-1}$ | Phytagel-based media were added to small agar plates in two distinct layers, to avoid settling of all carbon substrate at the bottom of plates;<br>During medium preparation, phytagel must be added very slowly to stirring medium to avoid lump formation |

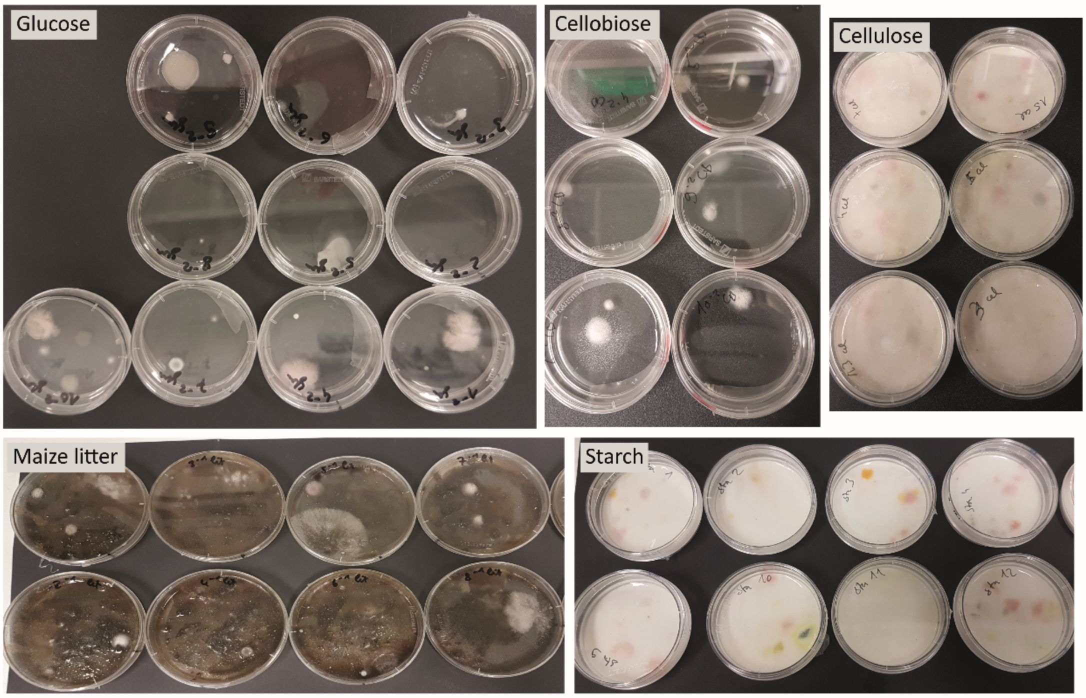


**Fig. S1** Soil dilution plating on different carbon substrate media to isolate diverse saprobic fungi. Examples of fungal isolation plates with different soil dilutions (ranging from full strength to $10^{-2}$ dilutions) after 3-6 days of growth are displayed. Individual colonies could well be differentiated and picked for isolation and individualization. Glucose and cellobiose media are solid phytagel-based growth media, while cellulose, starch and maize litter were added as sterile powder, supplemented with liquid media (Table S2). The addition of antibiotics successfully suppressed bacterial growth.

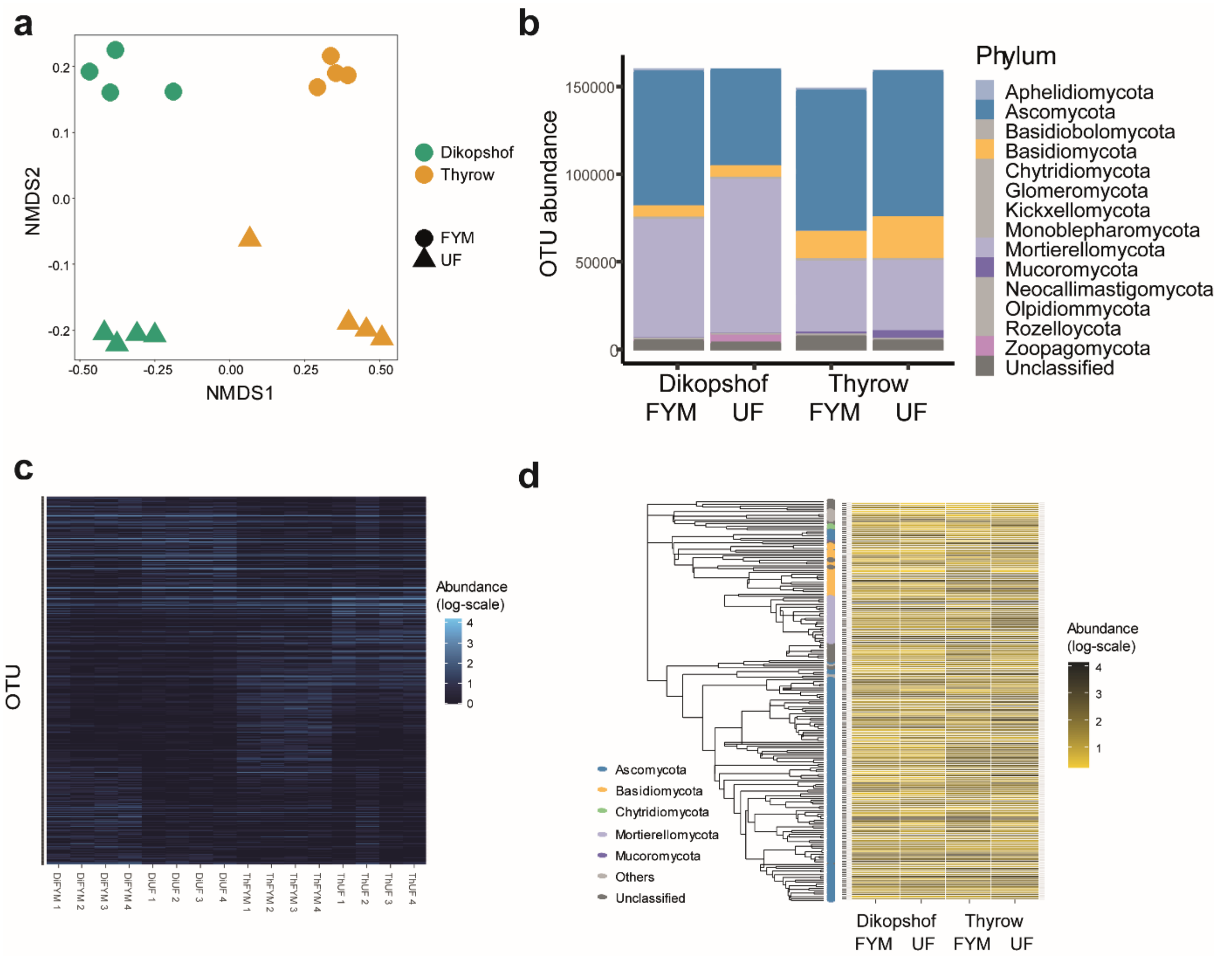


**Fig. S2**: Soil fungal community characteristics of sampled soil types for fungal isolation. Both locations, and respective fertilization treatments (Table S1) differ in community composition (**a**), phyla distribution (**b**) and taxa composition (**c, d**). **a** Multidimensional scaling of ASV abundance data of fungal communities (rarefied to a minimum of 40000 reads per sample based on rarefaction curves (vegan; Oksanen et al. 2022)). Colors differentiate soil types, shapes respective fertilization treatments. **b** Abundances of ASVs belonging to fungal phyla (differentiated by colors) in normalized ASV abundance tables. **c** Heatmap of OTU distribution in respective soil types, with light colors reflecting abundant OTUs (log10 scale). Taxa are sorted by their respective sample affiliation. **d** Heatmap of 200 most abundant OTUs in all samples, sorted by their phylogenetic placement (Neighbor-Joining tree (phangorn; Schliep 2010)). Colored dots at the tree tips indicate phyla affiliation, light to dark yellow colors of the heatmap respective average abundances in each soil type (log 10 scale).

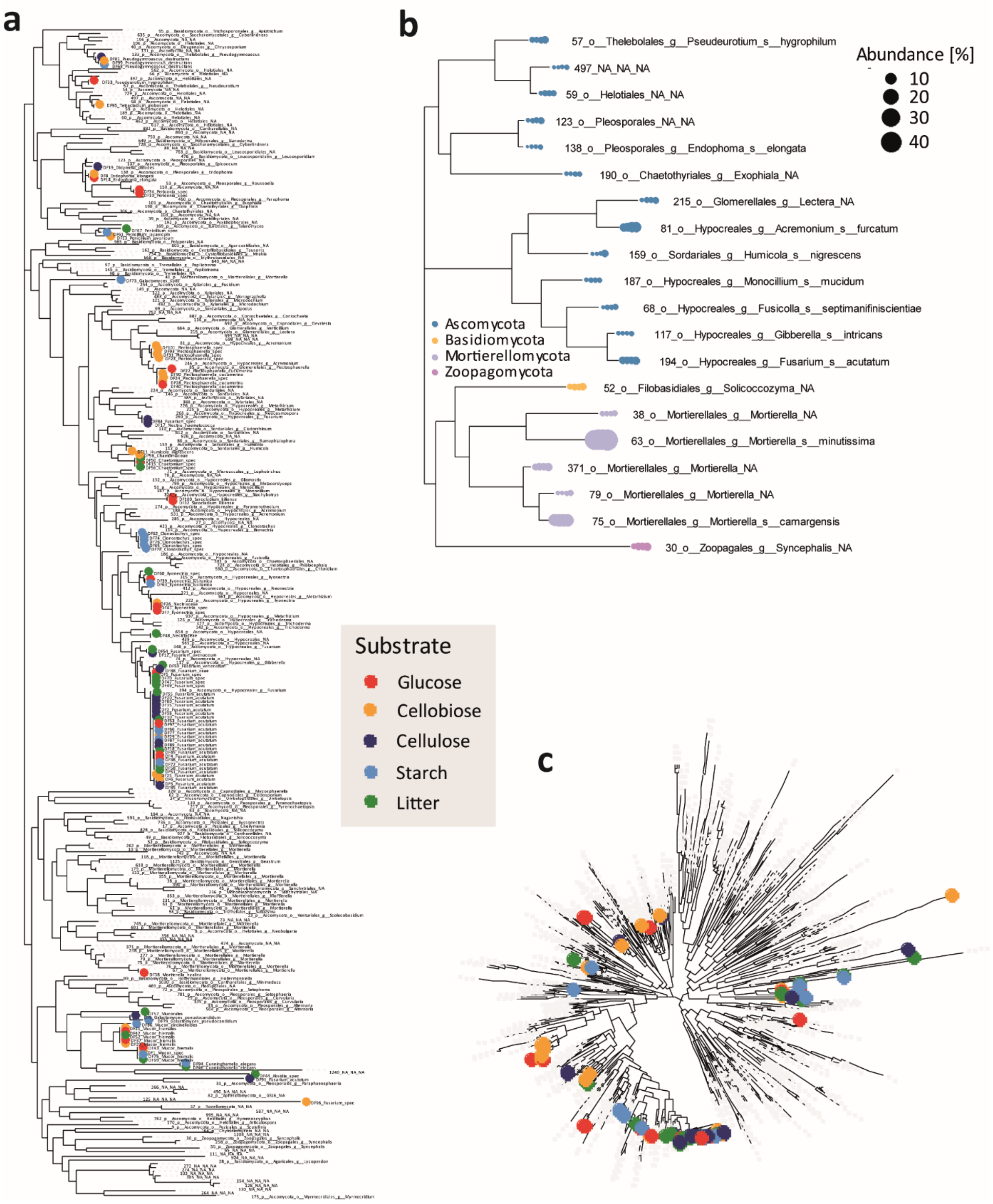


**Fig. S3**: Phylogenetic placement of isolated fungi in comparison to soil fungal community sequences, exemplified for Dikopshof soils with farmyard manure treatment. Fungal isolate sequences were reduced to ITS1 regions and aligned with Illumina sequences from respective soil types (Neighbor-Joining tree, *phangorn*). Colored dots illustrate fungal isolates differentiated by the carbon substrate medium they were isolated on (see legend). **a** The phylogenetic tree visualizes positions of fungal isolates in relation to the 204 most abundant OTUs observed in soil sequences, providing phylum, order and genus names (if available) as tip labels. Isolates are labelled with respective isolate IDs and species affiliation determined by long sequence reads (SSU, ITS and LSU regions). **b** Phylogenetic tree of the 20 most abundant species observed in soil communities (dot sizes indicate relative abundances in four replicate samples, colors refer to respective phyla). **c** The radial tree displays phylogenetic placement of fungal isolates in relation to the 500 most abundant fungal taxa in soil. Criteria to select fungal isolates (see Table S3 for final selection) were based on close proximity or overlap with abundant taxa in soil and an even representation of carbon sources used for isolation.

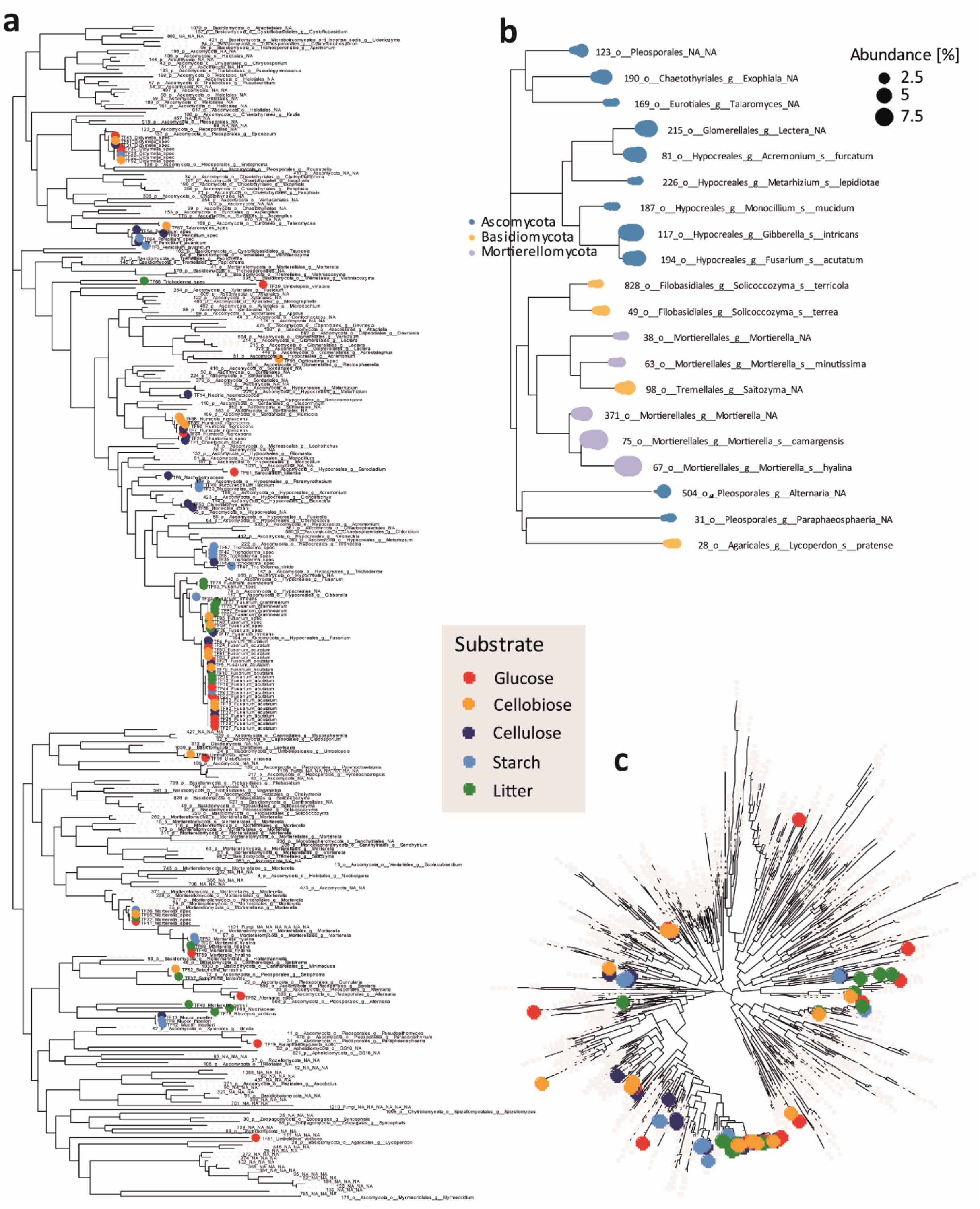


**Fig. S4**: Phylogenetic placement of isolated fungi in comparison to soil fungal community sequences, exemplified for Thyrow soils with farmyard manure treatment. Fungal isolate sequences were reduced to ITS1 regions and aligned with Illumina sequences from respective soil types (Neighbor-Joining tree, *phangorn*). Colored dots illustrate fungal isolates differentiated by the carbon substrate medium they were isolated on (see legend). **a** The tree visualizes the 204 most abundant taxa observed in soil sequences, providing phylum, order and genus names if available as tip labels. Isolates are labelled with respective isolate IDs and species affiliation determined by long sequence reads (ITS and LSU regions). **b** Phylogenetic tree of the 20 most abundant species observed in soil communities (dot sizes indicate relative abundances in four replicate samples, colors refer to respective phyla). **c** The radial tree displays phylogenetic placement of fungal isolates in relation to the 500 most abundant fungal taxa in soil. Criteria to select fungal isolates (see Table S3 for final selection) were based on close proximity or overlap with abundant taxa in soil and an even representation of carbon sources used for isolation.

**Table S3** Fungal isolates selected for further analyses based on their FunGuild classification and proximity to abundant soil taxa (Fig. S1; fungal identity determined based on long sequences of SSU, ITS and LSU regions compared to the Unite database (Nilsson et al. 2018) and RDP LSU dataset (Cole et al. 2014).

| **Isolate ID** | **Soil type** | **Phylum** | **Order** | **Species** | **Original carbon substrate used for isolation** | **FunGuild classification** | **NCBI accession numbers** | **DSMZ accession number** |
|---|---|---|---|---|---|---|---|---|
| DiF10 | Dik FYM | Ascomycota | Pleosporales | *Periconia_spec* | glucose | Pathotroph-Saprotroph-Symbiotroph | | |
| DiF11 | Dik FYM | Ascomycota | Sordariales | *Humicola_nigrescens* | cellobiose | Pathotroph-Saprotroph-Symbiotroph | | |
| DiF17 | Dik FYM | Ascomycota | Hypocreales | *Nectria_haematococca* | cellulose | Pathotroph-Saprotroph-Symbiotroph | | |
| DiF19 | Dik FYM | Ascomycota | Pleosporales | *Didymella_pinodes* | cellulose | Pathotroph-Saprotroph | | |
| DiF28 | Dik FYM | Ascomycota | Glomerellales | *Plectosphaerella_cucumeri* | cellobiose | Pathotroph-Symbiotroph | | |
| DiF33 | Dik FYM | Ascomycota | Thelebolales | *Pseudeurotium_hygrophilum* | glucose | Saprotroph | | |
| DiF34 | Dik FYM | Ascomycota | Pleosporales | *Periconia_spec* | glucose | Pathotroph-Saprotroph-Symbiotroph | | |
| DiF35 | Dik FYM | Ascomycota | Hypocreales | *Fusarium_acutatum* | cellulose | Pathotroph-Saprotroph-Symbiotroph | | |
| DiF38 | Dik FYM | Mortierellomycota | Mortierellales | *Mortierella_hyali* | glucose | Saprotroph-Symbiotroph | | |
| DiF41 | Dik FYM | Ascomycota | Hypocreales | *Ilyonectria_spec* | glucose | Pathotroph-Saprotroph-Symbiotroph | | |
| DiF44 | Dik FYM | Mucoromycota | Mucorales | *Absidia_spec* | litter | Saprotroph | | |

| | | | | | | |
|---|---|---|---|---|---|---|
| DiF56 | Dik FYM | Ascomycota | Sordariales | *Chaetomium_spec* | litter | Pathotroph-Saprotroph-Symbiotroph |
| DiF57 | Dik FYM | Mucoromycota | Mucorales | *Mucorales* | litter | Saprotroph |
| DiF59 | Dik FYM | Ascomycota | Hypocreales | *Fusarium_venetum* | litter | Pathotroph-Saprotroph-Symbiotroph |
| DiF60 | Dik FYM | Ascomycota | Hypocreales | *Ilyonectria_spec* | litter | Pathotroph-Saprotroph-Symbiotroph |
| DiF61 | Dik FYM | Ascomycota | Eurotiales | *Penicillium_javanicum* | starch | Saprotroph |
| DiF63 | Dik FYM | Ascomycota | Hypocreales | *Ilyonectria_lusitanica* | starch | Pathotroph-Saprotroph-Symbiotroph |
| DiF67 | Dik FYM | Ascomycota | Eurotiales | *Penicillium_spec* | litter | Saprotroph |
| DiF75 | Dik FYM | Mucoromycota | Mucorales | *Mucor_hiemalis* | starch | Saprotroph |
| DiF76 | Dik FYM | Ascomycota | Hypocreales | *Clonostachys_spec* | starch | Pathotroph-Saprotroph |
| DiF78 | Dik FYM | Ascomycota | Hypocreales | *Clonostachys_spec* | starch | Pathotroph-Saprotroph |
| DiF8 | Dik FYM | Ascomycota | Pleosporales | *Endophoma_elongata* | cellobiose | Pathotroph-Saprotroph |
| DiF81 | Dik FYM | Ascomycota | Thelebolales | *Pseudogymnoascus_destructans* | cellulose | Saprotroph |
| DiF84 | Dik FYM | Ascomycota | Hypocreales | *Fusarium_spec* | cellulose | Pathotroph-Saprotroph-Symbiotroph |
| DiF88 | Dik FYM | Ascomycota | Hypocreales | *Fusarium_zeae* | cellulose | Pathotroph-Saprotroph-Symbiotroph |
| DiF94 | Dik FYM | Mucoromycota | Mucorales | *Cunninghamella_elegans* | starch | Saprotroph |
| DiF95 | Dik FYM | Ascomycota | Helotiales | *Tetracladium_globosum* | cellobiose | Saprotroph-Symbiotroph |

| | | | | | | |
|---|---|---|---|---|---|---|
| DiF98 | Dik FYM | Ascomycota | Sordariales | *Chaetomiaceae* | cellobiose | Pathotroph-Saprotroph-Symbiotroph |
| ThF12 | Th FYM | Mucoromycota | Mucorales | *Mucor_moelleri* | starch | Saprotroph |
| ThF14 | Th FYM | Ascomycota | Hypocreales | *Nectria_haematococca* | cellulose | Pathotroph-Saprotroph-Symbiotroph |
| ThF19 | Th FYM | Ascomycota | Pleosporales | *Paraphaeosphaeria_spec* | glucose | Pathotroph-Saprotroph-Symbiotroph |
| ThF26 | Th FYM | Ascomycota | Sordariales | *Chaetomium_spec* | cellulose | Pathotroph-Saprotroph-Symbiotroph |
| ThF27 | Th FYM | Ascomycota | Hypocreales | *Fusarium_acutatum* | glucose | Pathotroph-Saprotroph-Symbiotroph |
| ThF28 | Th FYM | Ascomycota | Pleosporales | *Didymella_spec* | glucose | Pathotroph-Saprotroph |
| ThF3 | Th FYM | Ascomycota | Eurotiales | *Penicillium_javanicum* | starch | Saprotroph |
| ThF33 | Th FYM | Ascomycota | Hypocreales | *Fusarium_intricans* | starch | Pathotroph-Saprotroph-Symbiotroph |
| ThF35 | Th FYM | Ascomycota | Hypocreales | *Trichoderma_spec* | starch | Saprotroph |
| ThF36 | Th FYM | Mortierellomycota | Mortierellales | *Mortierella_spec* | starch | Saprotroph-Symbiotroph |
| ThF37 | Th FYM | Ascomycota | Pleosporales | *Setophoma_terrestris* | litter | Pathotroph-Saprotroph |
| ThF40 | Th FYM | Ascomycota | Hypocreales | *Purpureocillium_lilacinum* | starch | Pathotroph-Symbiotroph |
| ThF49 | Th FYM | Mortierellomycota | Mortierellales | *Mortierella_gamsii* | litter | Saprotroph-Symbiotroph |
| ThF51 | Th FYM | Mucoromycota | Umbelopsidales | *Umbelopsis_vicea* | glucose | Saprotroph |

| | | | | | | |
|---|---|---|---|---|---|---|
| ThF53 | Th FYM | Ascomycota | Hypocreales | *Clonostachys_spec* | cellulose | Pathotroph-Saprotroph |
| ThF54 | Th FYM | Ascomycota | Hypocreales | *Trichoderma_spec* | cellulose | Saprotroph |
| ThF58 | Th FYM | Ascomycota | Hypocreales | *Bionectria_solani* | cellulose | Pathotroph-Saprotroph |
| ThF59 | Th FYM | Mortierellomycota | Mortierellales | *Mortierella_hyali* | glucose | Saprotroph-Symbiotroph |
| ThF6 | Th FYM | Ascomycota | Hypocreales | *Stachybotryaceae* | cellulose | Saprotroph |
| ThF62 | Th FYM | Ascomycota | Pleosporales | *Alterria_spec* | glucose | Pathotroph-Saprotroph |
| ThF65 | Th FYM | Ascomycota | Hypocreales | *Fusarium_graminearum* | litter | Pathotroph-Saprotroph-Symbiotroph |
| ThF72 | Th FYM | Mortierellomycota | Mortierellales | *Mortierella_spec* | litter | Saprotroph-Symbiotroph |
| ThF74 | Th FYM | Ascomycota | Hypocreales | *Fusarium_aveceum* | litter | Pathotroph-Saprotroph-Symbiotroph |
| ThF81 | Th FYM | Ascomycota | Pleosporales | *Didymella_spec* | cellobiose | Pathotroph-Saprotroph |
| ThF82 | Th FYM | Ascomycota | Pleosporales | *Setophoma_terrestris* | cellobiose | Pathotroph-Saprotroph |
| ThF84 | Th FYM | Mucoromycota | Umbelopsidales | *Umbelopsis_spec* | cellobiose | Saprotroph |
| ThF86 | Th FYM | Ascomycota | Sordariales | *Humicola_nigrescens* | cellobiose | Pathotroph-Saprotroph-Symbiotroph |
| ThF87 | Th FYM | Ascomycota | Eurotiales | *Talaromyces_spec* | cellobiose | Saprotroph |
| ThF93 | Th FYM | Ascomycota | Ophiostomatales | *Ophiostoma_spec* | cellobiose | Pathotroph-Saprotroph |
| DiU1 | Dik UF | Ascomycota | Hypocreales | *Metarhizium_marquandii* | cellobiose | Pathotroph-Symbiotroph |
| DiU12 | Dik UF | Ascomycota | Sordariales | *Schizothecium_curvisporum* | starch | Saprotroph |

| | | | | | | |
|---|---|---|---|---|---|---|
| DiU15 | Dik UF | Ascomycota | Hypocreales | *Dactylonectria_estremocensis* | cellobiose | Pathotroph-Saprotroph-Symbiotroph |
| DiU16 | Dik UF | Ascomycota | Hypocreales | *Neonectria_lugdunensis* | cellobiose | Pathotroph-Saprotroph-Symbiotroph |
| DiU19 | Dik UF | Ascomycota | Pleosporales | *Periconia_spec* | cellobiose | Pathotroph-Saprotroph-Symbiotroph |
| DiU21 | Dik UF | Ascomycota | Glomerellales | *Plectosphaerella_cucumeri* | cellobiose | Pathotroph-Symbiotroph |
| DiU23 | Dik UF | Ascomycota | Microascales | *Cephalotrichum_stemonitis* | cellobiose | Pathotroph-Saprotroph-Symbiotroph |
| DiU25 | Dik UF | Ascomycota | Xylariales | *Microdochium_spec* | cellobiose | Pathotroph-Symbiotroph |
| DiU35 | Dik UF | Ascomycota | Hypocreales | *Ilyonectria_macrodidyma* | glucose | Pathotroph-Saprotroph-Symbiotroph |
| DiU40 | Dik UF | Ascomycota | Hypocreales | *Fusarium_aveceum* | litter | Pathotroph-Saprotroph-Symbiotroph |
| DiU42 | Dik UF | Ascomycota | Pleosporales | *Curvularia_fallax* | glucose | Pathotroph-Saprotroph |
| DiU45 | Dik UF | Basidiomycota | Cantharellales | *Minimedusa_spec* | glucose | Pathotroph |
| DiU49 | Dik UF | Ascomycota | Glomerellales | *Plectosphaerella_spec* | glucose | Pathotroph-Saprotroph-Symbiotroph |
| DiU51 | Dik UF | Ascomycota | Chaetothyriales | *Exophiala_equi* | glucose | Pathotroph-Saprotroph |
| DiU52 | Dik UF | Ascomycota | Xylariales | *Xylariales* | glucose | NA |
| DiU58 | Dik UF | Ascomycota | Hypocreales | *Trichoderma_hamatum* | cellulose | Saprotroph |
| DiU6 | Dik UF | Mortierellomycota | Mortierellales | *Mortierella_spec* | starch | Saprotroph-Symbiotroph |

| | | | | | | |
|---|---|---|---|---|---|---|
| DiU63 | Dik UF | Ascomycota | Hypocreales | *Fusarium_intricans* | litter | Pathotroph-Saprotroph-Symbiotroph |
| DiU64 | Dik UF | Ascomycota | Pleosporales | *Didymella_pinodes* | litter | Pathotroph-Saprotroph |
| DiU68 | Dik UF | Ascomycota | Thelebolales | Pseudogymnoascus_destructans | cellulose | Saprotroph |
| DiU7 | Dik UF | Ascomycota | Hypocreales | Fusarium_acutatum | litter | Pathotroph-Saprotroph-Symbiotroph |
| DiU74 | Dik UF | Ascomycota | Hypocreales | Clonostachys_spec | cellulose | Pathotroph-Saprotroph |
| DiU78 | Dik UF | Ascomycota | Hypocreales | Trichoderma_spec | cellulose | Saprotroph |
| DiU86 | Dik UF | Ascomycota | Hypocreales | Fusarium_spec | cellulose | Pathotroph-Saprotroph-Symbiotroph |
| ThU10 | Th UF | Ascomycota | Hypocreales | Fusarium_spec | cellulose | Pathotroph-Saprotroph-Symbiotroph |
| ThU14 | Th UF | Ascomycota | Sordariales | Chaetomiaceae | cellulose | Pathotroph-Saprotroph-Symbiotroph |
| ThU17 | Th UF | Ascomycota | Venturiales | Rhizosphaera_pini | glucose | Pathotroph-Saprotroph-Symbiotroph |
| ThU19 | Th UF | Ascomycota | Pleosporales | Didymella_spec | cellobiose | Pathotroph-Saprotroph |
| ThU21 | Th UF | Ascomycota | Capnodiales | Cladosporium_delicatulum | litter | Pathotroph-Saprotroph-Symbiotroph |
| ThU22 | Th UF | Ascomycota | Hypocreales | Fusarium_gibbosum | litter | Pathotroph-Saprotroph-Symbiotroph |
| ThU26 | Th UF | Ascomycota | Glomerellales | Plectosphaerella_spec | cellobiose | Pathotroph-Saprotroph-Symbiotroph |

| | | | | | | |
|---|---|---|---|---|---|---|
| ThU30 | Th UF | Ascomycota | Hypocreales | Nectriaceae | litter | Pathotroph-Saprotroph-Symbiotroph |
| ThU33 | Th UF | Ascomycota | Hypocreales | Fusarium_spec | litter | Pathotroph-Saprotroph-Symbiotroph |
| ThU36 | Th UF | Ascomycota | Hypocreales | Fusarium_acutatum | cellulose | Pathotroph-Saprotroph-Symbiotroph |
| ThU37 | Th UF | Ascomycota | Capnodiales | Cladosporium_spec | litter | Pathotroph-Saprotroph-Symbiotroph |
| ThU38 | Th UF | Ascomycota | Sordariales | Chaetomiaceae | cellulose | Pathotroph-Saprotroph-Symbiotroph |
| ThU4 | Th UF | Ascomycota | Eurotiales | Penicillium_spec | cellulose | Saprotroph |
| ThU42 | Th UF | Ascomycota | Sordariales | Arcopilus_cupreus | starch | Pathotroph-Saprotroph-Symbiotroph |
| ThU47 | Th UF | Mucoromycota | Mucorales | Mucor_moelleri | litter | Saprotroph |
| ThU49 | Th UF | Ascomycota | Hypocreales | Trichoderma_spec | starch | Saprotroph |
| ThU58 | Th UF | Ascomycota | Eurotiales | Talaromyces_spec | cellobiose | Saprotroph |
| ThU64 | Th UF | Mortierellomycota | Mortierellales | Mortierella_gemmifera | cellobiose | Saprotroph-Symbiotroph |
| ThU72 | Th UF | Ascomycota | Ophiostomatales | Ophiostoma_spec | glucose | Pathotroph |
| ThU75 | Th UF | Mucoromycota | Umbelopsidales | Umbelopsis_spec | glucose | Saprotroph |
| ThU78 | Th UF | Ascomycota | Thelebolales | Pseudogymnoascus_roseus | starch | Saprotroph |
| ThU81 | Th UF | Ascomycota | Hypocreales | Fusarium_acutatum | starch | Pathotroph-Saprotroph-Symbiotroph |

| | | | | | | |
|---|---|---|---|---|---|---|
| ThU9 | Th UF | Ascomycota | Pleosporales | Epicoccum_dendrobii | litter | Pathotroph-Saprotroph |
| ThU92 | Th UF | Ascomycota | Hypocreales | Niesslia_spec | glucose | Saprotroph |

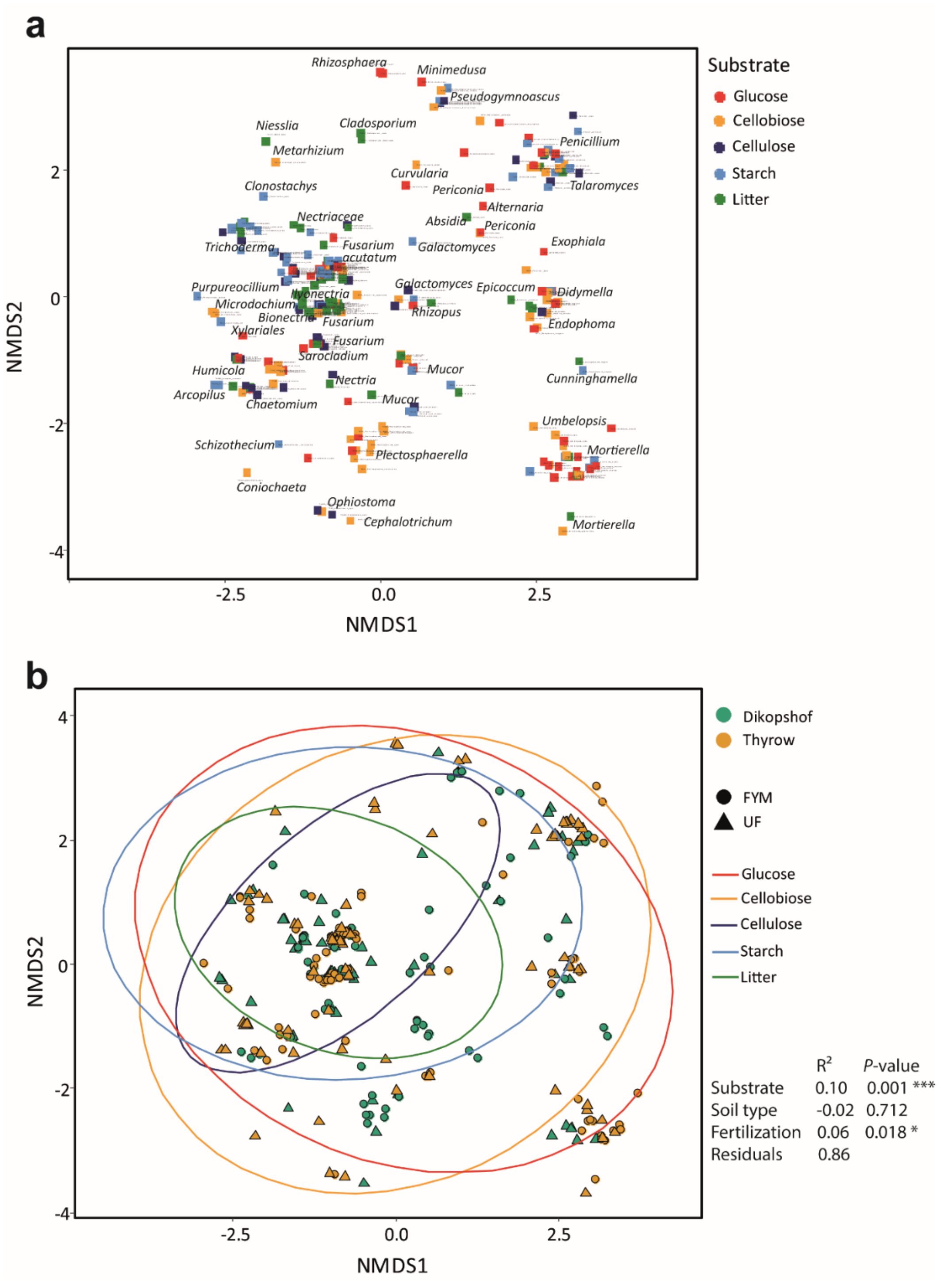


**Fig. S5**: Correlation of phylogenetic distance among all fungal isolates in relation to carbon substrates used for isolation and soil origin. Phylogenetic distance was calculated based on alignment of long sequence reads (SSU, LSU and iTS region) with MAFFT (ips; Heibl 2008), visualizing pairwise distances (*phangorn*) with non-metric multidimensional scaling (NMDS, *vegan*). **a** Individual dots represent fungal isolates, labelled with ID and species/genus identification. Colors indicate respective carbon substrate isolation media (see legend). The location of different genera is indicated to provide phylogenetic information. **b** The impact of original soil type and fertilization treatment is visualized by dot colors and shapes. Ellipses visualize the 95% confidence region of carbon substrates used for isolation. The table displays statistical results of substrate, soil type and soil fertilization effects on the phylogenetic distance in fungal isolates, analyzed by permutational multivariate analysis of variance (*vegan*).

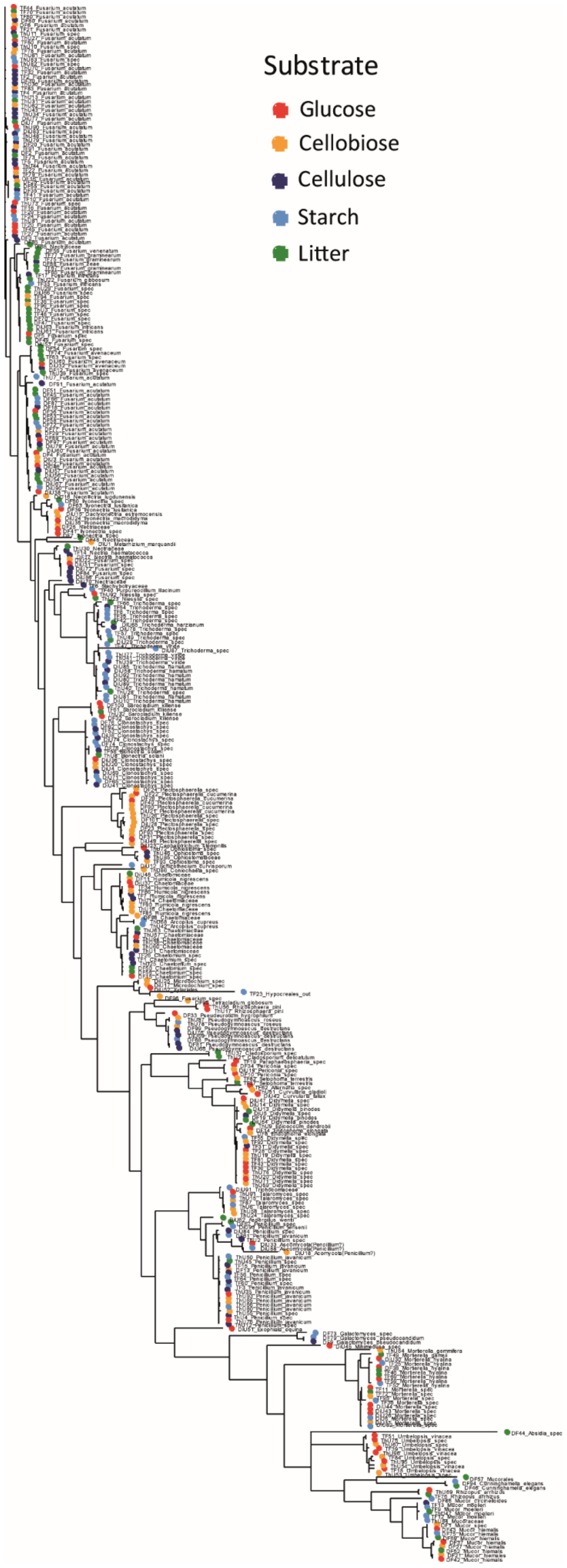


**Fig. S6**: Phylogenetic tree displaying all fungal isolates obtained in this study, isolated on different carbon substrates (visualized by respective colored dots, see legend). The Neighbor-joining tree (*phangorn*) was built based on an alignment of long sequence reds (SSU, ITS, LSU region) with MAFFT (*ips*).

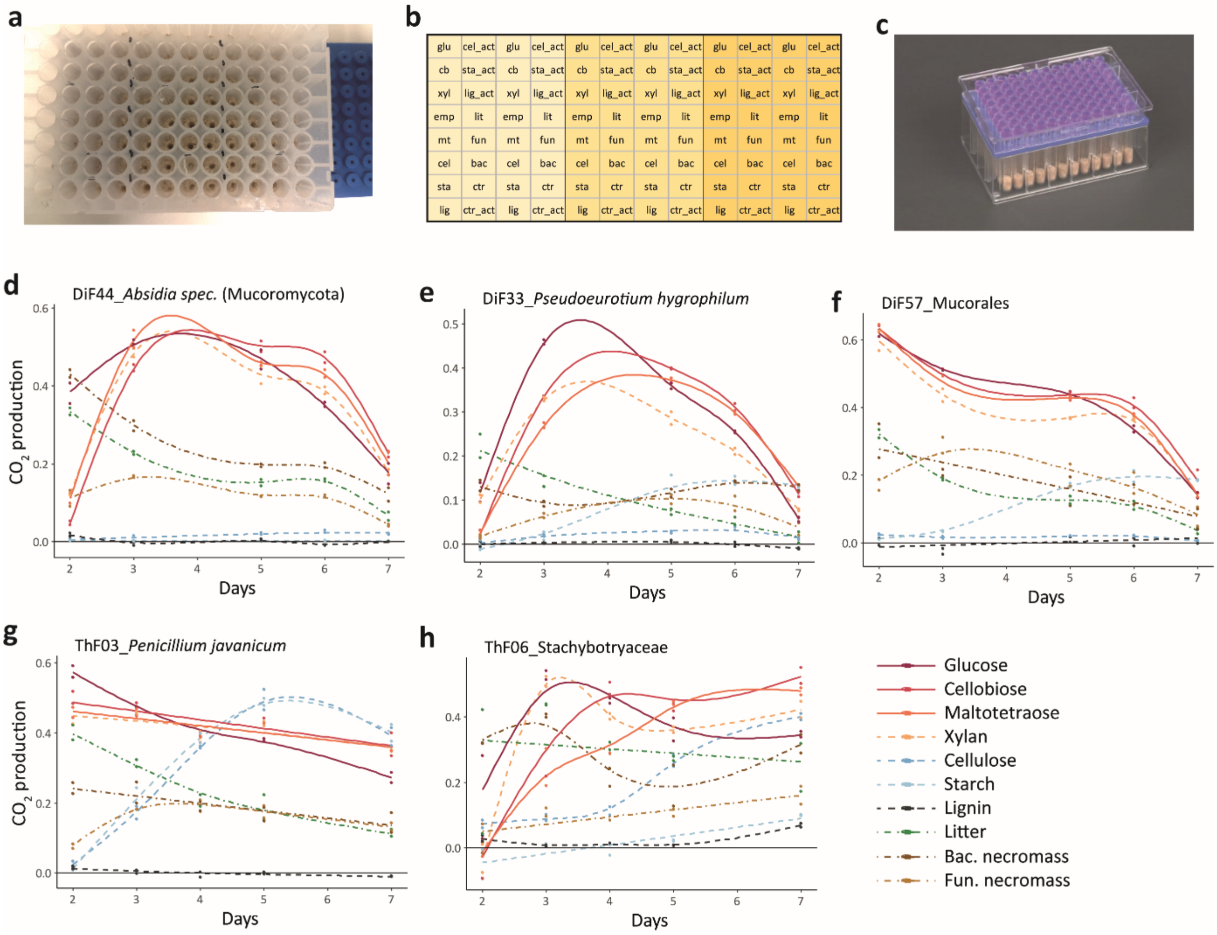


**Fig. S7**: Experimental design of respiration measurements of the novel *FungiResp* approach, testing for fungal activity on different carbon substrates. **a-c** Experimental design of 96-well plates, displaying the inoculation of fungi into artificial soil media with different carbon substrates (**a**), arrangement of substrates and replicates/isolates within plates, different colors indicate the placement of individual isolates and replicates (**b**) and the MicroResp$^{TM}$ detection plates (**c**; image taken from https://www.microresp.com/). **d-h** Curves displaying generalized additive models (gam, *mgcv* (Wood 2011)) fitted to respiration measurements over time measured by the *FungiResp* approach, displayed for exemplary fungal isolates and treatments (colours and line shapes visualize carbon substrates added (see legend), dots represent individual data points). For absolute respiration activity over time (Fig. 3), the area under the curve was calculated by integral functions.

**Table S4** Defined liquid media used in main trait experiments with the *FungiResp* approach.

| Element | Compound | Medium concentration | Comments | Manufacturer |
|---|---|---|---|---|
| C | Glucose $C_6H_{12}O_6$ | 5.00 g $L^{-1}$ | Carbon sources were added at equal molar amounts, calculations were based on molecular formulas (when given) or total C contents (CN analyzer (EuroEA, HekaTech, Germany)); Activated complex carbon (_act) includes 5% glucose, calculated as % of total molar C needed; Carbon sources were autoclaved separately to avoid medium caramelization; pH of carbon substrate solutions was adjusted to pH of glucose (~5.6) | |
| | Cellobiose $C_{12}H_{22}O_{11}$ | 4.75 g $L^{-1}$ | | D-(+)-Cellobiose, 98%, AB114600 (abcr GmbH, Karlsruhe, Germany) |
| | Maltotetraose $C_{24}H_{42}O_{21}$ | 4.62 g $L^{-1}$ | | Maltotetraose, >90%, O-MAL4 (Megazyme Neogen®, Bray, Ireland) |
| | Xylan $(C_5H_8O_4)n$ | 4.40 g $L^{-1}$ | | Xylan, ≥95 % Xylooligosaccharides from corncob, 8659.1 (Carl Roth GmbH + Co. KG, Karlsruhe, Germany) |
| | Cellulose $(C_6H_{10}O_5)n$ | 4.50 g $L^{-1}$ | | Cellulose, Type 101, S6790 (Merck KGaA, Darmstadt, Germany) |
| | Cellulose_act | 4.27 g $L^{-1}$ + 0.22 g $L^{-1}$ glucose | | |
| | Starch (C6H10O5)n | 4.5 g $L^{-1}$ | | Starch from corn, S4126 (Merck KGaA, Darmstadt, Germany) |
| | Starch_act | 4.27 g $L^{-1}$ + 0.22 g $L^{-1}$ glucose | | |
| | Lignin | 3.17 g $L^{-1}$ | | Lignin, alkali, 370959 (Merck KGaA, Darmstadt, Germany) |
| | Lignin_act | 3.01 g $L^{-1}$ + 0.22 g $L^{-1}$ glucose | | |
| | Maize litter | 4.37 g $L^{-1}$ | | |
| | Fungal necromass | 3.60 g $L^{-1}$ | | |
| | Bacterial necromass | 4.04 g $L^{-1}$ | | |
| | Control | | | |
| N | $NH_4NO_3$ | 0.503 g $L^{-1}$ | | |
| P | $NaH_2PO_4$ | 0.183 g $L^{-1}$ | Phosphate solution was autoclaved separately to avoid insoluble precipitates or toxic conditions in the presence of salts (Tanaka et al. 2014) | |
| Mg, S | $MgSO_4$ | 0.067 g $L^{-1}$ | | |
| K | KCl | 0.023 g $L^{-1}$ | | |
| Ca | $CaCl_2$ | 0.065 g $L^{-1}$ | | |
| Fe | NaFeEDTA | 0.02 g $L^{-1}$ | | |
| vitamine | Thiamine HCl | 1 mg $L^{-1}$ | Micronutrients and vitamins were added from stock solutions | |
| vitamine | Biotin | 0.05 mg $L^{-1}$ | | |
| Mo | $Na_2MoO_4$ | 0.05 mg $L^{-1}$ | | |
| Cu | $CuSO_4$ | 0.01 mg $L^{-1}$ | | |

| | | |
|---|---|---|
| **Mn** | $MnSO_4$ | 0.05 mg $L^{-1}$ |
| **B** | $H_3BO_3$ | 0.05 mg $L^{-1}$ |
| **Zn** | $ZnSO_4$ | 5 mg $L^{-1}$ |

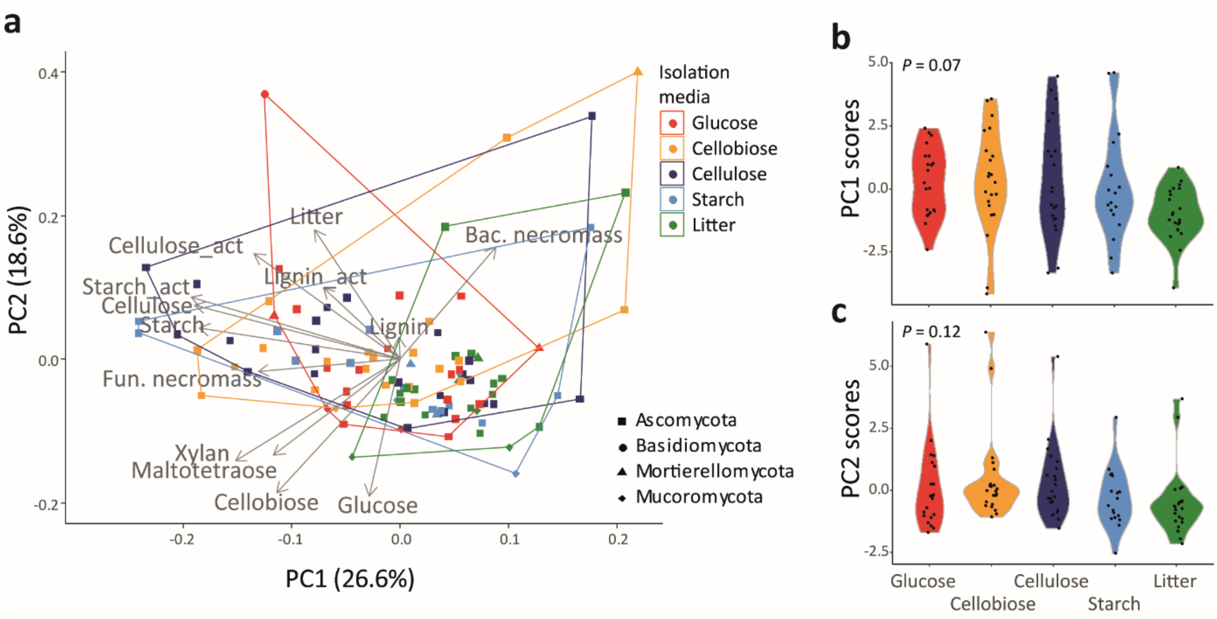


**Fig. S8**: Correlation of carbon substrate use ability of individual isolates with the original media used for isolation. **a** Principle component analysis (PCA) of carbon substrate use ability of complex C sources (Fig. 5a), marking fungal isolates by color with respective isolation media used (see legend). **b, c** PC scores of isolates on PC1 (**b**) and PC2 (**c**) in relation to isolation media used based on respective carbon sources. *P*-values are based on one-way analyses of variances, testing for overall isolation substrate effects.

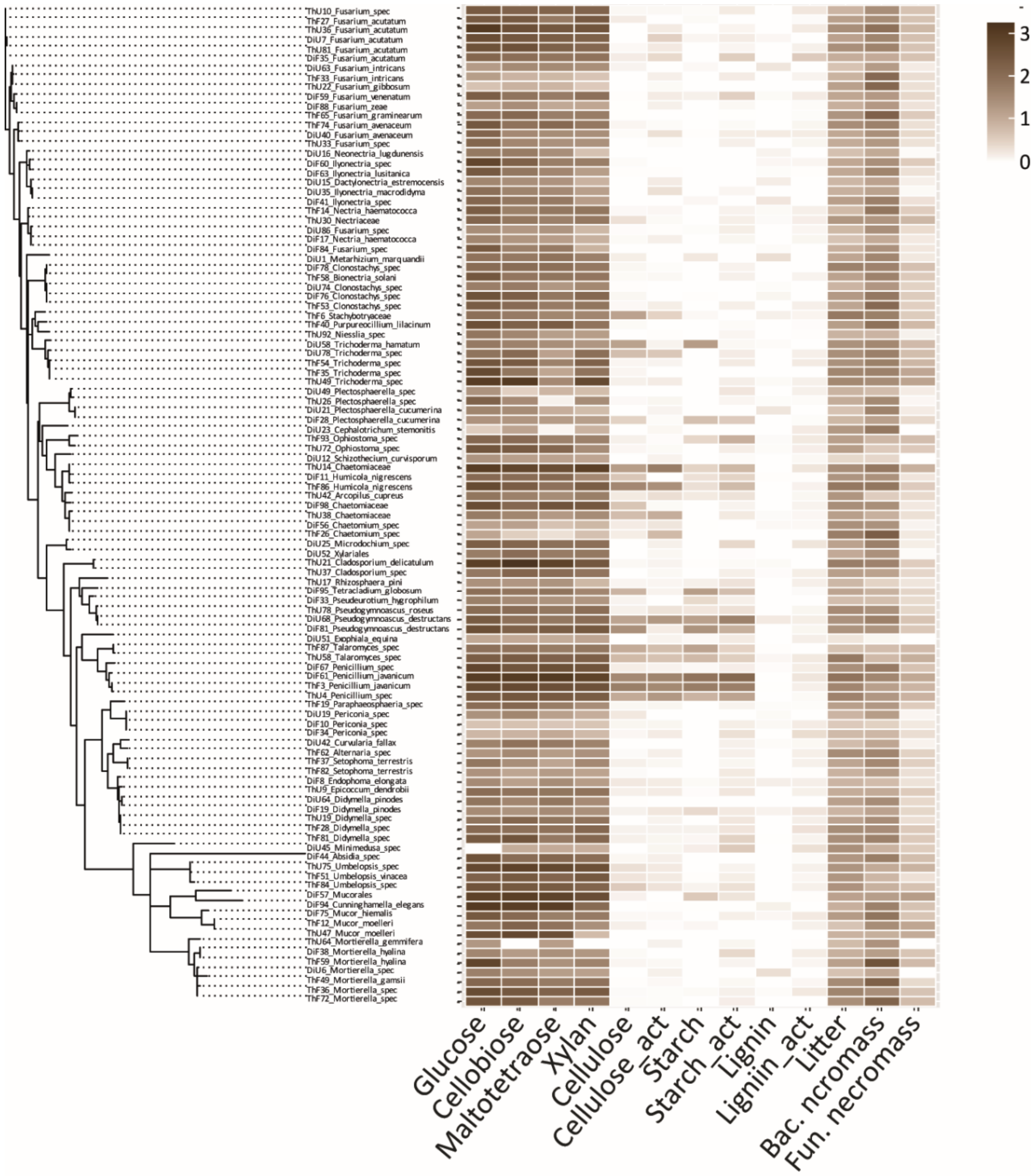


**Fig. S9**: Carbon substrate use ability in different fungal isolates, sorted by phylogenetic placement with individual tip labels added (Fig. 3). The heatmap visualizes fungal activity on carbon substrates varying in complexity, given as $CO_2$ production over 7 days (n = 2; values represent integral functions over time, based on MicroResp values subtracted by the control (no C added) of respective isolates). Isolates are sorted based on their phylogenetic placement (neighbor-joining tree; Fig. 3).

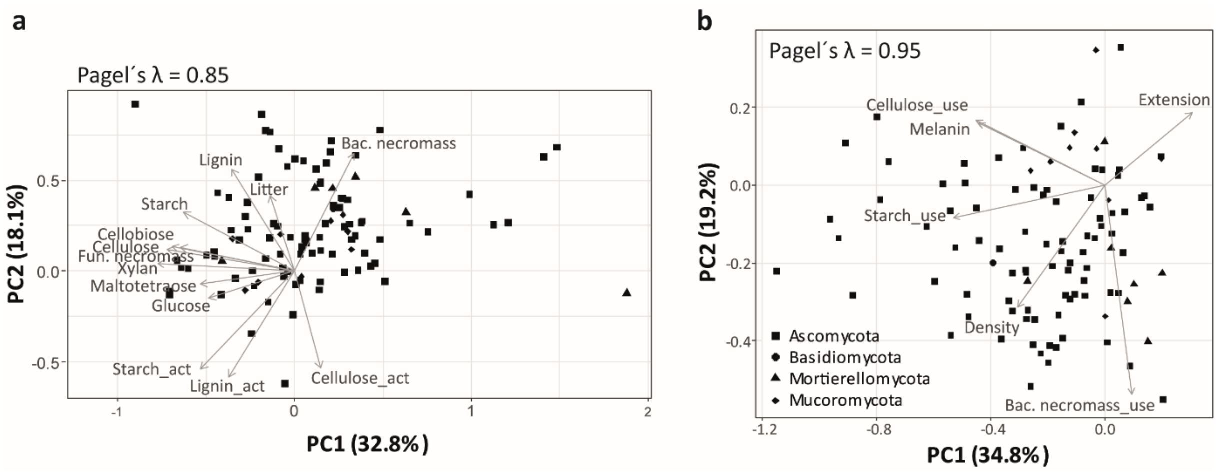


**Fig. S10**: Phylogenetic principal component analyses (PCA) displaying correlations of fungal isolate ability to use various complex carbon substrates (**a**) and its relation to the fungal economics spectrum (**b**). We used phylogenetic PCA to analyze correlation structure independent of evolutionary history (phytools; Revell 2012). The first two principal component (PC) axes are displayed, including respective percentages of variability explained. Arrows represent eigenvectors of traits on PC axes, dots individual isolates (shapes of dots indicate phylogenetic placement).

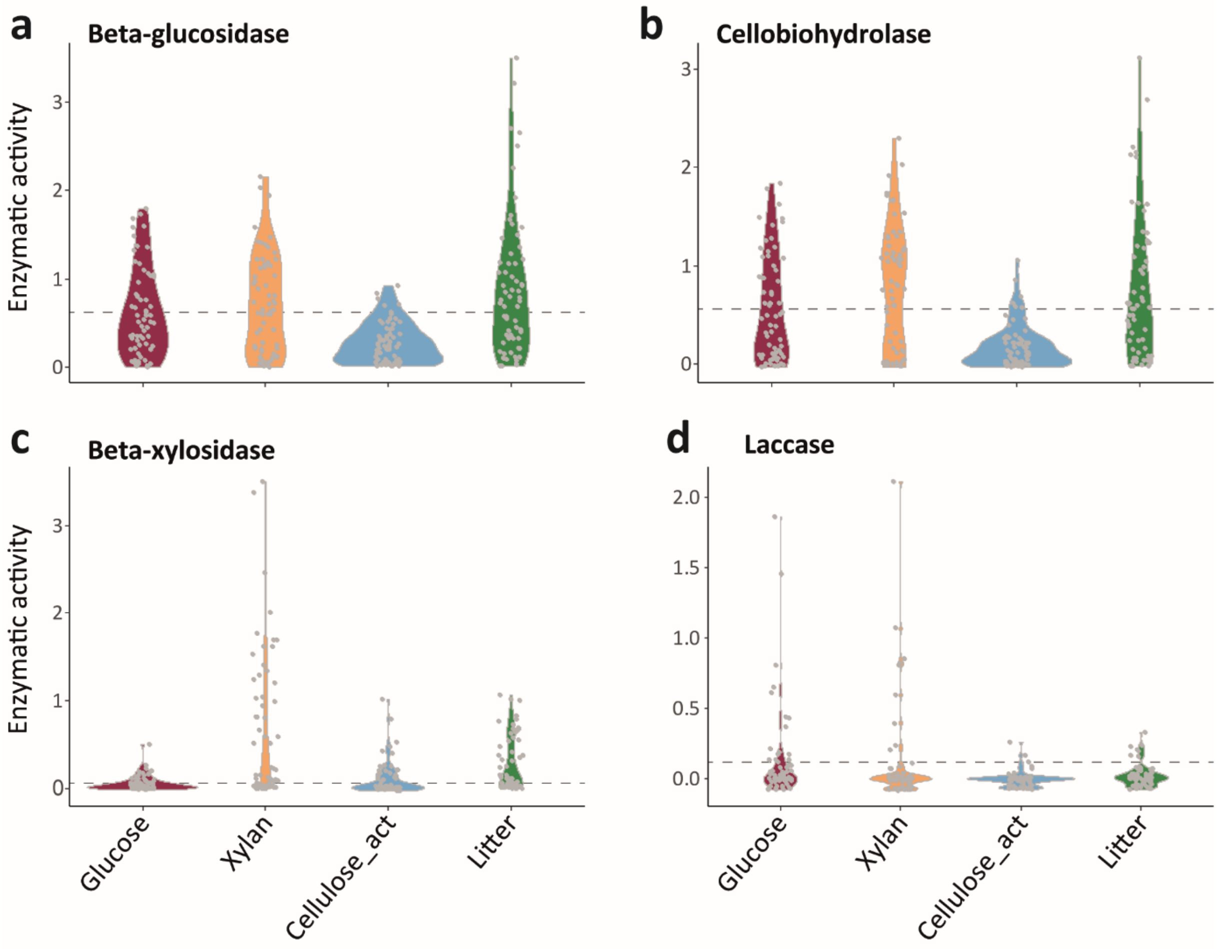

**Fig. S11**: Average enzymatic activity of fungal isolates grown on different carbon source media, based on colorimetric assays testing for beta-glucosidase (**a**), cellobiodhydrolase (**b**), beta-xylosidase (**c**) and laccase (**d**). Data distribution is illustrated by violin plots, individual data points are jittered on the plot (32 isolates, n=2). Dashed lines indicate mean enzymatic activity of respective enzymes on glucose media.

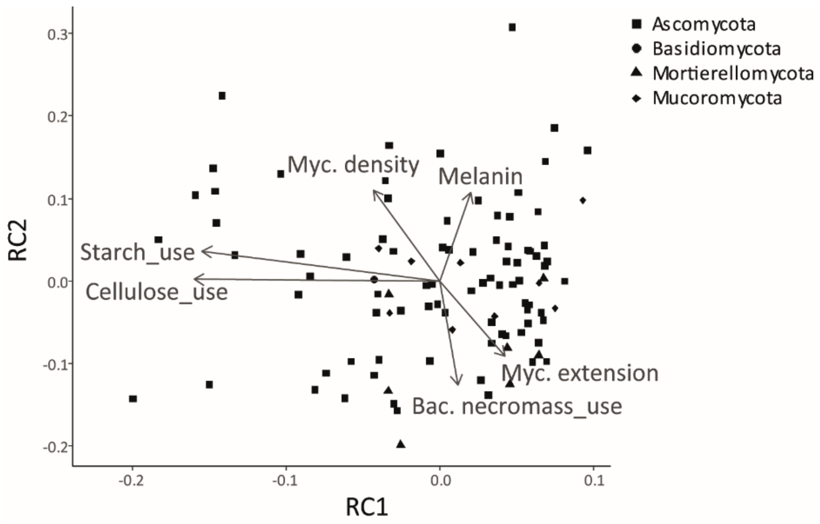


**Fig. S12**: Varimax rotated principal component analysis of complex carbon use ability with fungal traits displaying the fungal economics spectrum (Fig. 4b). The first two rotated components (RC) are displayed. Arrows represent eigenvectors of traits on RC axes, dots individual isolates (shapes of dots refer to phylogenetic placement). Varimax rotation allows the rotation of PC axes in order to maximize trait loadings on resulting rotated components (RC), while maintaining the orthogonal (linearly uncorrelated) structure among different axes (Camenzind et al. 2024).

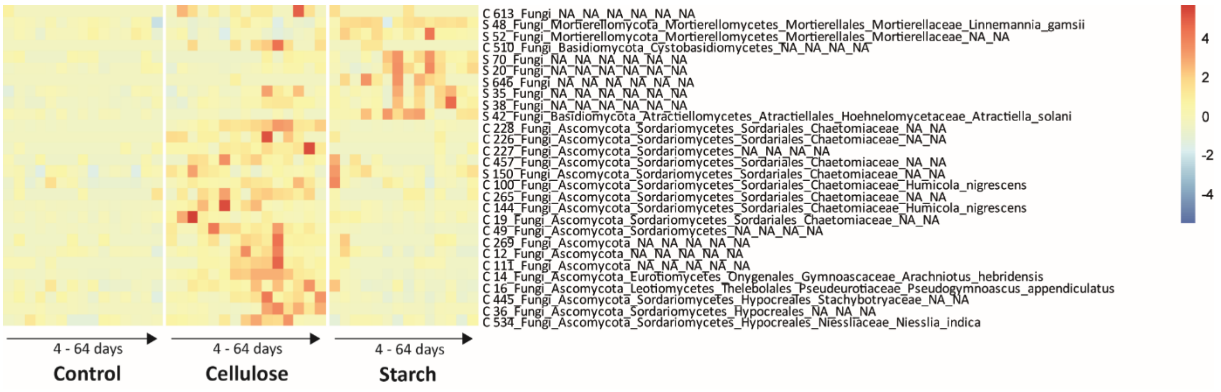


**Fig. S13:** Heatmap of fungal OTUs significantly increased by substrate additions to soil (MaAslin 2 results; Mallick et al. 2021). Colors indicate OTU abundances in soil communities, based on Hellinger transformed community data. Only OTUs significantly increased in individual treatments are displayed (qval < 0.05 (FDR-adjusted *P*-value), including time as random factor). OUT names are based on UNITE classification of ITS1 sequences (Nilsson et al. 2018), letters indicate significant positive correlations with respective treatments (S: starch, C: cellulose).

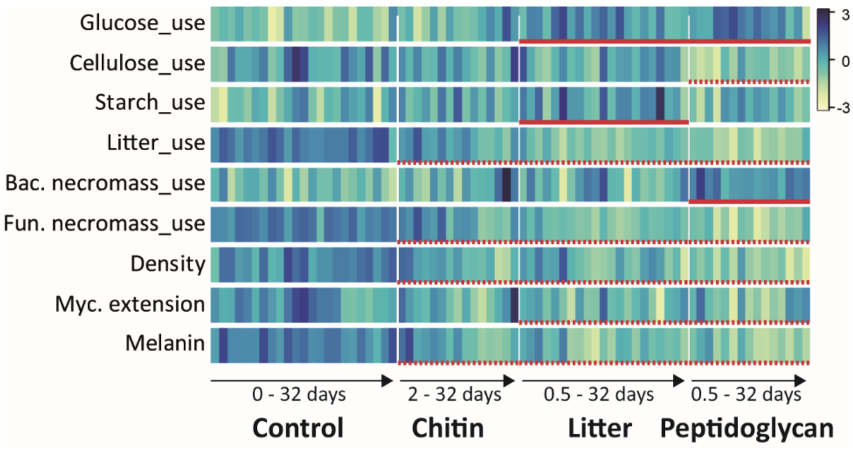


**Fig. S14:** Functional soil fungal community shifts in response to chitin, litter and peptidoglycan additions (Vonhoegen et al., in prep.). Shifts in Community-level weighted means of individual functional traits are visualized as heatmap, displaying functional shifts among treatments over time (3 replicates per time point). Darker colour indicates high relative values of scaled community-level weighted means (mean = 0, standard deviation = 1). Significant difference to the control are indicated by red lines, dashed lines represent negative effects.

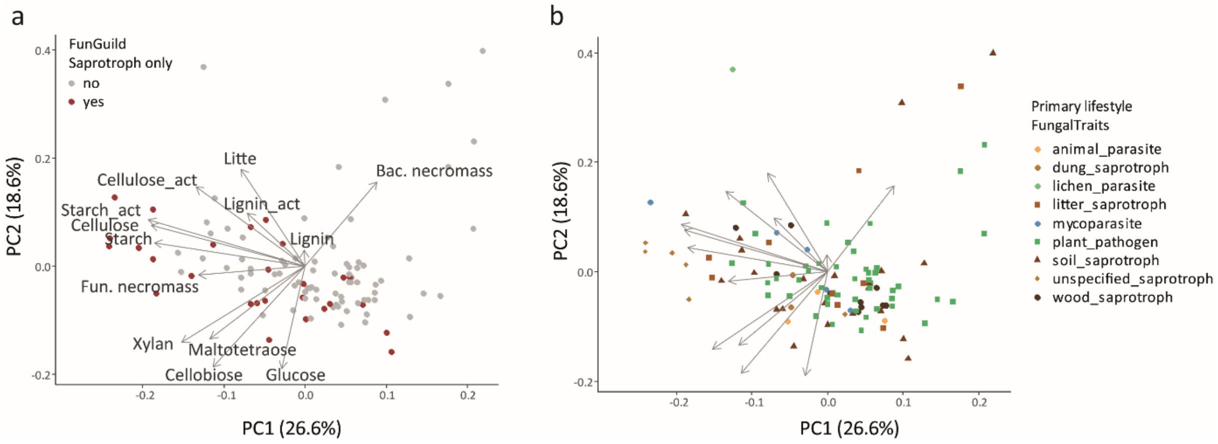


**Fig. S15**: Classification of fungal isolates by guilds according to FunGuild (**a**) and FungalTraits (**b**), sorted within the trait space related to carbon use ability traits (Nguyen et al. 2016, Põlme et al. 2020). **a** Isolates classified as Saprotroph only are marked in brown, isolates classified as Saprotroph as well as Symbiotroph and/or Pathotroph are marked in grey. **b** The primary lifestyle of fungal isolates as defined by the FungalTraits database is indicated by respective colors and shapes. For details on principal component analyses please refer to Fig. 4a.

Põlme, S., K. Abarenkov, R. Henrik Nilsson, B. D. Lindahl, K. E. Clemmensen, H. Kauserud, N. Nguyen, R. Kjøller, S. T. Bates, P. Baldrian, T. G. Frøslev, K. Adojaan, A. Vizzini, A. Suija, D. Pfister, H.-O. Baral, H. Järv, H. Madrid, J. Nordén, J.-K. Liu, J. Pawlowska, K. Põldmaa, K. Pärtel, K. Runnel, K. Hansen, K.-H. Larsson, K. D. Hyde, M. Sandoval-Denis, M. E. Smith, M. Toome-Heller, N. N. Wijayawardene, N. Menolli, N. K. Reynolds, R. Drenkhan, S. S. N. Maharachchikumbura, T. B. Gibertoni, T. Læssøe, W. Davis, Y. Tokarev, A. Corrales, A. M. Soares, A. Agan, A. R. Machado, A. Argüelles-Moyao, A. Detheridge, A. de Meiras-Ottoni, A. Verbeken, A. K. Dutta, B.-K. Cui, C. K. Pradeep, C. Marín, D. Stanton, D. Gohar, D. N. Wanasinghe, E. Otsing, F. Aslani, G. W. Griffith, T. H. Lumbsch, H.-P. Grossart, H. Masigol, I. Timling, I. Hiiesalu, J. Oja, J. Y. Kupagme, J. Geml, J. Alvarez-Manjarrez, K. Ilves, K. Loit, K. Adamson, K. Nara, K. Küngas, K. Rojas-Jimenez, K. Bitenieks, L. Irinyi, L. G. Nagy, L. Soonvald, L.-W. Zhou, L. Wagner, M. C. Aime, M. Öpik, M. I. Mujica, M. Metsoja, M. Ryberg, M. Vasar, M. Murata, M. P. Nelsen, M. Cleary, M. C. Samarakoon, M. Doilom, M. Bahram, N. Hagh-Doust, O. Dulya, P. Johnston, P. Kohout, Q. Chen, Q. Tian, R. Nandi, R. Amiri, R. H. Perera, R. dos Santos Chikowski, R. L. Mendes-Alvarenga, R. Garibay-Orijel, R. Gielen, R. Phookamsak, R. S. Jayawardena, S. Rahimlou, S. C. Karunarathna, S. Tibpromma, S. P. Brown, S.-K. Sepp, S. Mundra, Z.-H. Luo, T. Bose, T. Vahter, T. Netherway, T. Yang, T. May, T. Varga, W. Li, V. R. M. Coimbra, V. R. T. de Oliveira, V. X. de Lima, V. S. Mikryukov, Y. Lu, Y. Matsuda, Y. Miyamoto, U. Kõljalg, and L. Tedersoo. 2020. FungalTraits: a user-friendly traits database of fungi and fungus-like stramenopiles. Fungal Diversity 105:1-16.

Revell, L. J. 2012. phytools: an R package for phylogenetic comparative biology (and other things). Methods in Ecology and Evolution 3:217-223.

Schliep, K. P. 2010. phangorn: phylogenetic analysis in R. Bioinformatics 27:592-593.

Tanaka, T., K. Kawasaki, S. Daimon, W. Kitagawa, K. Yamamoto, H. Tamaki, M. Tanaka, C. H. Nakatsu, and Y. Kamagata. 2014. A hidden pitfall in the preparation of agar media undermines microorganism cultivability. Applied and Environmental Microbiology 80:7659-7666.

Wood, S. N. 2011. Fast stable restricted maximum likelihood and marginal likelihood estimation of semiparametric generalized linear models. Journal of the Royal Statistical Society: Series B (Statistical Methodology) 73:3-36.
